# Distinct basal ganglia contributions to learning from implicit and explicit value signals in perceptual decision-making

**DOI:** 10.1101/2023.09.05.556317

**Authors:** Tarryn Balsdon, M. Andrea Pisauro, Marios G. Philiastides

## Abstract

Metacognitive evaluations of confidence provide an estimate of decision accuracy that could guide learning in the absence of explicit feedback. We examine how humans might learn from this implicit feedback in direct comparison with that of explicit feedback, using simultaneous EEG-fMRI. Participants performed a motion direction discrimination task where stimulus difficulty was increased to maintain performance, with intermixed explicit- and no-feedback trials. We isolate single-trial estimates of post-decision confidence using EEG decoding, and find these neural signatures re-emerge at the time of feedback together with separable signatures of explicit feedback. We identified these signatures of implicit versus explicit feedback along a dorsal-ventral gradient in the striatum, a finding uniquely enabled by an EEG-fMRI fusion. These two signals are then integrated into an aggregate representation in the external globus pallidus, which broadcasts updates to improve cortical decision processing via the thalamus and insular cortex, irrespective of the source of feedback.

## Introduction

Practicing a task helps to improve performance [1]. In tasks involving a behavioural response to an external stimulus, performance can benefit from learning at the level of action selection, decision processes, or indeed, sensitivity in perceiving the task-relevant features of the stimulus. Explicit feedback about whether responses were correct has been shown to increase learning rate at each of these levels [2,3]. These performance improvements, however, have also been shown to occur in the absence of explicit feedback [1,3,4]. Though several variables could contribute to improved performance in the absence of explicit feedback (including mere exposure to a stimulus [5]), one variable of particular interest is the metacognitive evaluation of decision confidence. Confidence provides an estimate of the likelihood that a decision is correct [6,7], and as such could be used as an implicit proxy for explicit feedback.

Our understanding of the role of confidence in learning lags that of explicit feedback. The influence of explicit feedback on learning can be understood from a reinforcement learning framework, which traditionally described how one learns to select actions that maximise future external rewards in value-based learning environments [8]. More recently this framework has also been successfully applied to perceptual learning [2]. In this context, reinforcement-like learning can occur based on explicit feedback information without external rewards [9] or purely based on the observer’s internal estimate of confidence as a proxy for feedback [10,11]. This suggests that the computational description of learning behaviour could be generalised to incorporate different feedback signals [12]. However, it is unclear whether the same neural processes used for explicit feedback are flexibly appropriated to implement learning from implicit signals, such as confidence.

To date, there has been no direct comparison of the neural mechanisms for learning from confidence to those of learning from explicit feedback within the same experimental task. Ideally, this comparison would be made across intermixed trials where explicit feedback is periodically available, as in ecological contexts [13]. However, in these contexts it could be beneficial to wait for infrequent but reliable feedback than implement learning based on confidence estimates that are an imprecise estimate of decision accuracy [14,15] or that might be vulnerable to biases [16–18]. Alternatively, confidence could provide a more fine-grained estimate of performance, reflecting the graded precision of underlying decision processes [19], as opposed to the binary outcome of explicit feedback (which is also blind to the processing that led to the outcome). In this way, confidence could be valuable in providing more nuanced information of how to adjust decision-making processes, even in the presence of explicit feedback. In the case where confidence has a proportionate impact even in the presence of explicit feedback, this might necessitate not only separate neural signatures for the different signals but also the presence of a downstream process in which the two signals form an aggregate reinforcement-like representation to jointly influence learning.

The dopaminergic system is well known for its role in organising reward-seeking behaviour and motor responses [20,21]. The striatum, in particular, has been repeatedly linked to the representation of expected reward [22], action values [23,24], reward prediction errors [25,26], and more intrinsic motivational (hedonistic) signals [27], including confidence [11,28,29]. Similarly, the presence of extensive cortico-striatal circuits [30] along with the cortico-basal ganglia-thalamocortical loop [31], which are amenable to plastic changes via phasic dopaminergic firing [32], make the basal ganglia well positioned for implementing learning updates and guiding future actions. However, it is of ongoing debate how distinct subregions within the basal ganglia might distinctly or jointly contribute to learning [33,34], and whether there is a distinction in the computation of explicit versus implicit feedback signals.

Here, we provide observers with intermittent feedback during a perceptual decision-making task to test whether observers learn by exploiting their internal confidence estimates, even when explicit feedback is frequently available. Patterns of behavioural perseveration suggest that this was indeed the case. We use this design of intermixed trials with and without explicit feedback to directly compare the neural signatures of learning from explicit (outcome value) and implicit (confidence) feedback signals. We first isolate endogenous electrophysiological trial-wise estimates of confidence and outcome value using a decoding analysis of EEG. We then harness these trial-wise estimates to inform the analysis of simultaneously acquired fMRI data. We traced the BOLD signatures of implicit and explicit feedback to a spatial gradient in the striatum, where implicit feedback is represented more dorsally while explicit feedback is represented more ventrally. Moreover, these two signals of implicit and explicit feedback appear to integrate into an aggregate representation in the external globus pallidus, which broadcasts updates to improve cortical decision processing via the thalamus, irrespective of whether the source of feedback included explicit feedback or not.

## Results

Participants (N = 23) performed a variant of the classic random dot motion direction discrimination task [35] (see **Methods**) whilst simultaneous EEG-fMRI was recorded. Each trial was composed of three time-windows (**Figure 1A**): The decision-window (in which the stimulus was presented and perceptual decision reported); the bet-window (in which participants were given the opportunity to bet that their responses were correct); and the feedback-window (in which participants were cued whether they would receive explicit feedback or not, and if so, shown the points awarded for the trial). On each trial, 1 point was gained for a correct response, or 1 point lost for an incorrect response. These values were doubled if the participant bet that they made the correct response. The total accumulated points corresponded to a monetary bonus at the end of the experiment. On explicit-feedback trials, participants were shown the points gained/lost on that trial, but on no-feedback trials participants could only infer how many points were awarded, and they were cued to do so. Explicit- and no-feedback trials were intermixed throughout the experiment.

**Figure 1.**
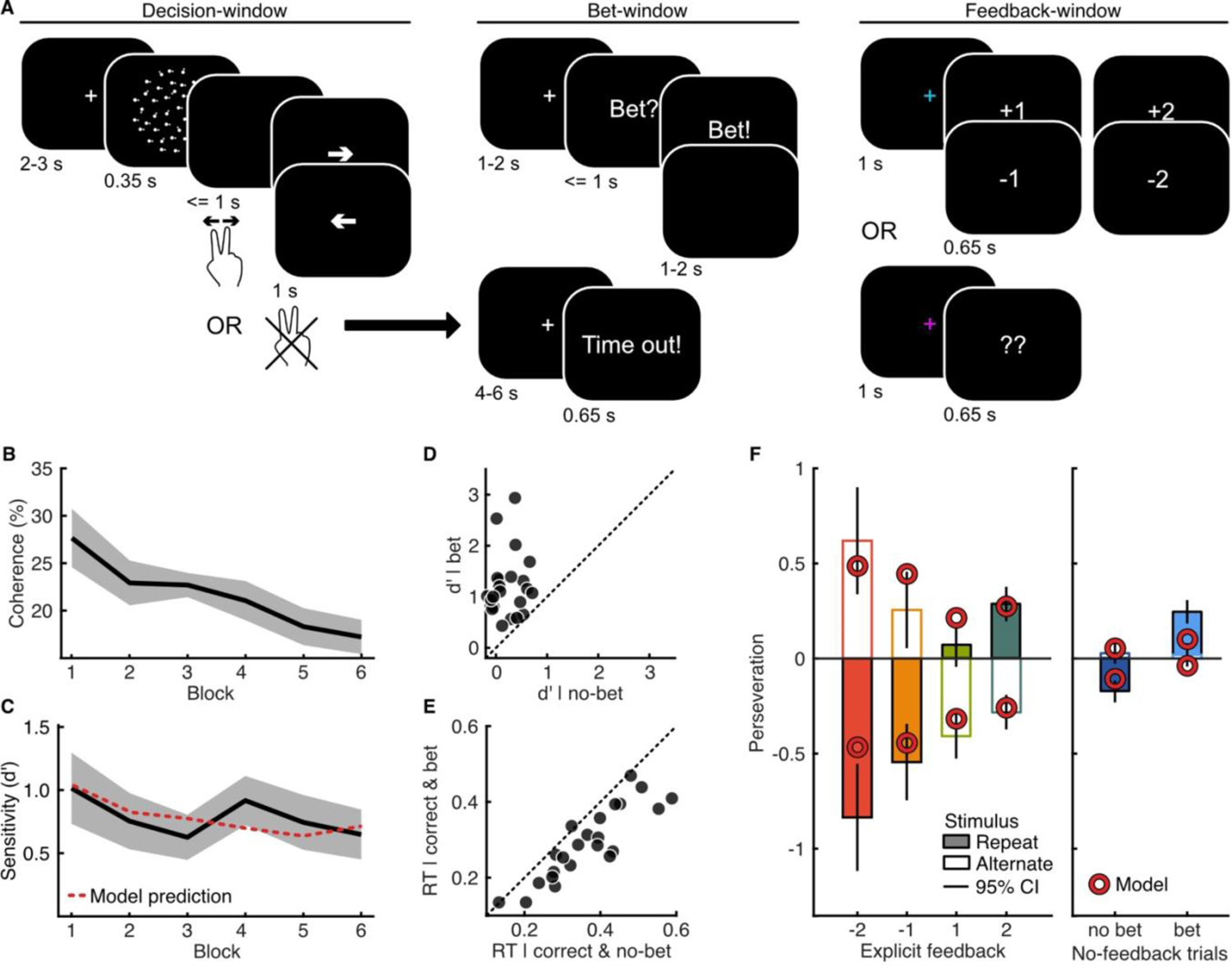
Methods and behaviour. **A)** Trials consisted of three time-windows: the decision-window; bet-window; and feedback-window. In the decision widow, a fixation cross was presented for a variable duration, followed by 350 ms of random dot motion, and up to 1 second to enter the left/right response, with visual feedback about the button pressed. In the bet-window, participants were given up to a second to bet that their response was correct, to double the value of the points gained/lost on that trial. In the feedback-window, participants were cued (coloured fixation) about whether they would receive explicit feedback, or be shown two question marks indicating they should think about how many points they think they won/lost on that trial. If participants failed to enter their left/right decision within one second after the stimulus, they were informed with the words ‘time out’, they lost a point, and the trial was excluded from the analysis. **B)** Coherence (percent of dots moving in the correct direction) was adjusted each block of 50 trials to maintain accuracy around 65% correct. Shaded region shows 95% within-subject confidence intervals (on the difference across blocks). **C)** Sensitivity (d’) across blocks, with 95% within-subject confidence intervals shaded. The dashed line shows the prediction of the simple computational model using confidence to learn. **D)** Sensitivity (d’) on bet (ordinate) and no-bet (abscissa) trials. **E)** Median reaction time (seconds) on correct trials for bet (ordinate) and no-bet (abscissa) trials. **F)** Perseveration, measured as the normalised probability of repeating a response, for repeat (filled) and alternate (open) stimuli, relative to the overall probability of response repetition, by feedback on the previous trial, or the previous bet response on no-feedback trials (error bars show 95% CI within-subjects difference between repeat and alternate stimuli). The markers show the predictions of the simple computational model. Note that perseveration for alternate stimuli was close to 0 for both bet and no-bet no-feedback trials, and so are barely visible, but present.

### Behavioural signatures of learning from confidence as implicit feedback

Task difficulty (proportion of coherent dots) was controlled across blocks (6 in total) to maintain performance between 55-75% correct (see **Methods**). There was an overall increase in task difficulty (**Figure 1B**, mean difference in coherence between first and last block = −10.43% ± 4.67, 95% within subject confidence interval, t(22) = 4.64, *p* < 0.001) with no substantial change in sensitivity (**Figure 1C**, mean difference in d’ = −0.37 ± 0.45 t(22) = 1.67, *p* = 0.109, with Bayes factor, calculated based on the savage-dickey ratio with a unit-information prior, BF_10_ = 0.75, representing insubstantial evidence in favour of the null hypothesis for no difference in sensitivity) suggesting participants learnt to improve at the task during the course of the experiment. We designed this experiment such that participants were frequently given explicit feedback (50% of randomly intermixed trials), and so did not have to rely on decision confidence as the sole source of information to improve their performance. Evidence for learning on no-feedback trials could therefore be considered robust evidence for the involvement of confidence in learning.

Although the overall tendency to bet may partly depend on inter-individual differences in risk aversion, there were also signs that bet responses did reflect a metacognitive evaluation of decision confidence: participants were more likely to be correct when they bet (difference in d’ = 0.98 ± 0.28; t(22) = 7.35, *p*<0.001; **Figure 1D**) and they also showed faster reaction times (mean difference in median reaction time = 0.07 s ± 0.02, t(22) = 6.63, *p* < 0.001; **Figure 1E**).

We found response perseveration (the interaction between response and stimulus repetition), depended on the sign of the previous explicit feedback, a typical signature of feedback-learning (**Figure 1F, left;** 2 (value, 1 vs 2) x 2 (sign, + vs -) repeated measures ANOVA, main effect of sign F(1,22) = 60.04, *p* < 0.001 after Bonferroni correction for three comparisons; non-significant main effect of value F(1,22) = 2.23, *p* = 0.15 uncorrected; and the interaction, F(1,22) = 5.18, *p* = 0.033, which would not survive Bonferroni correction). Given the design of intermixed explicit-feedback and no-feedback trials, participants could have relied solely on explicit feedback to improve their performance. However, we found the same pattern of response perseveration on no-feedback trials, depending on whether the participant bet on their previous response (**Figure 1F, right;** within-subjects t-test, t(22) = 4.69, *p* < 0.001). This is in line with the hypothesis that high confidence is treated as a positive outcome, to be used as reinforcement. Indeed, a simple computational reinforcement learning model suggested that using confidence as the expected value across all trials was a better description of the data (dashed line in **Figure 1C** and markers in **1F**) compared to a more traditional model with no learning on no-feedback trials (∑ Δ*BIC* = −330.45; exceedance probability > 0.99; see **Methods, Supplementary Figure 1**). These behavioural results are evidence that confidence is used as an implicit learning signal in the absence of explicit feedback.

### Neural signatures of implicit and explicit feedback

We used an asymmetric (EEG to fMRI) fusion analysis [36,37] of the simultaneous EEG-fMRI data. To examine the EEG signatures of confidence as implicit feedback, a Linear Discriminant Analysis (LDA, see **Methods**) was used to isolate weights on the EEG channel activity (spatial filter) that best discriminated bet from no-bet trials in the decision-window. The summed product of the spatial filter and channel activity gives the bet-prediction, a continuous variable where, for each trial, the higher the bet-prediction the more likely the trial was a bet trial. Spatial filters were first generated for each time-point within the decision-window, and then a robust individual filter was selected for each participant as the average of five consecutive filters that best discriminated bet from no-bet trials within the group-level significant time-window (see **Methods**). Applying these individual spatial filters over time within a trial shows how the EEG activity relevant for discriminating bet from no-bet trials emerges over time (**Figure 2A**). The bet-prediction not only discriminated bet from no-bet trials (from the time of the response to 0.25 s after, mean F(1,22) = 15.36, cluster corrected *p* < 0.001; **Figure 2A, left)**, but also showed a significant main effect of response accuracy (from 0.1 to 0.24 s after the response, mean F(1,22) = 6.66, cluster corrected *p* = 0.002), and predicted reaction times on correct trials (mean GLM β-weight = 0.15 ± 0.06, t(22) = 5.31, *p* < 0.001). In this way, the EEG bet-prediction could reflect a more graded representation of the underlying confidence used to arbitrate whether or not to bet on individual trials.

**Figure 2.**
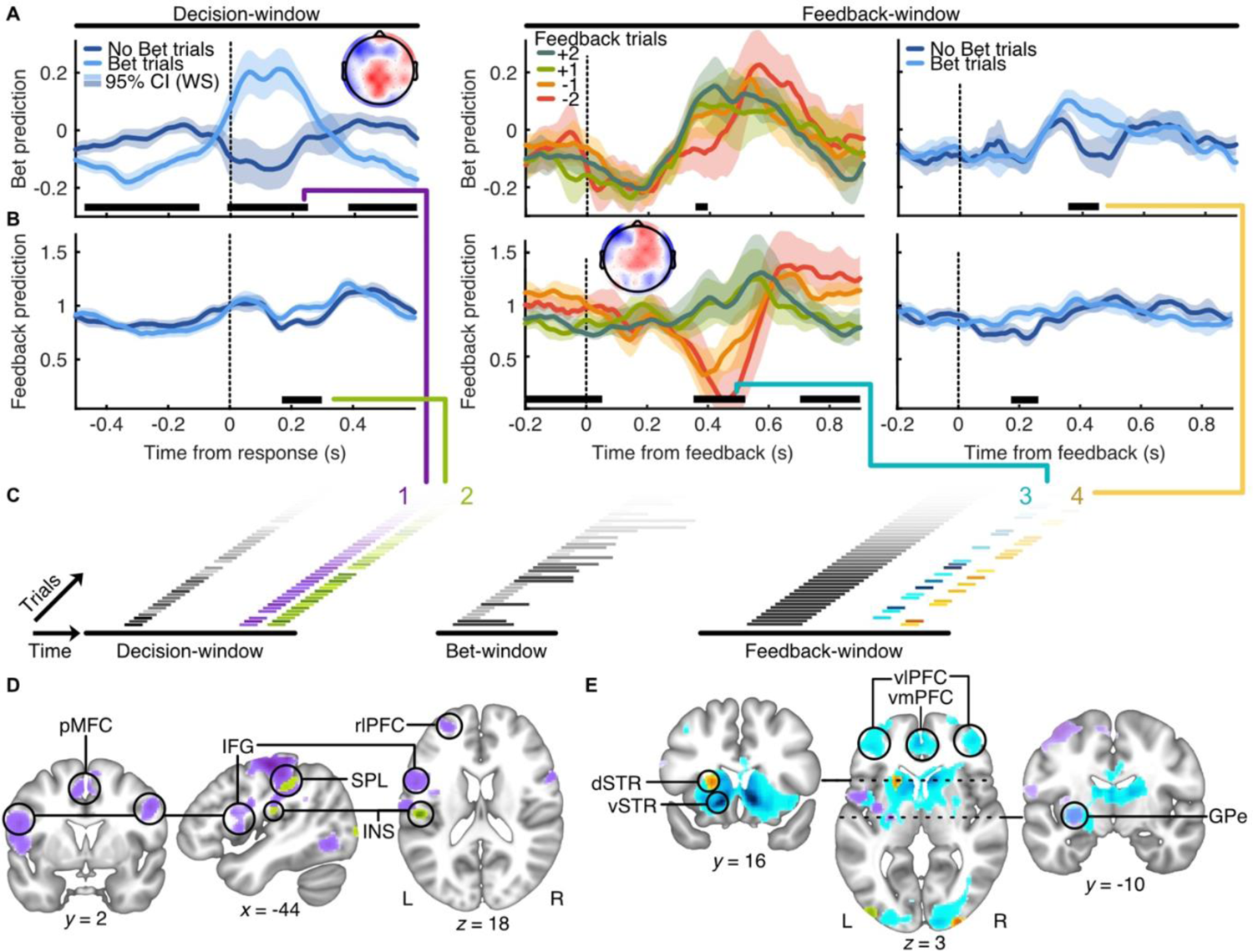
EEG signatures of implicit and explicit feedback and their BOLD correlates. **A)** EEG bet-prediction in the decision-window (left) and the feedback-window for explicit feedback (middle) and no-feedback (right) trials. The bet-prediction was generated by selecting the spatial filter that best discriminated bet from no-bet responses in the decision-window (insert shows the group average scalp projection of the discriminating components of this spatial filter). The spatial filter is applied to the EEG over time to generate a prediction for each trial, the lines show the average grouping trials by whether the participant bet, or the explicit feedback given. “Time from feedback” refers to the time from the explicit feedback or the cue to infer the points earnt (question marks) as shown in Figure 1A. Shaded regions indicate 95% within-subjects confidence intervals (on the difference across trial conditions), and black horizontal lines show windows of cluster-corrected significant differences. **B)** The same as A) for the feedback-prediction, generated by selecting the spatial filter (insert shows the group average scalp projection of the discriminating components) that best discriminated feedback-sign (positive vs. negative) during the feedback-window (middle). **C)** Construction of the EEG-informed fMRI GLM analysis with four EEG regressors (coloured) and traditional regressors (grey) which corresponded to stimulus onset (with amplitude modulated by decision time), bet-cue (with duration modulated by bet response time), and feedback cue. Time within a trial progresses left to right, with example trials layered. All regressors had a duration of 0.1s, except for the feedback cue (1 s, duration to feedback) and the bet-cue on bet-trials (black bars, extended to the duration of the bet response). **D)** Clusters of voxels with a significant positive relation to the EEG bet-prediction (1, purple) and feedback-prediction (2, green) in the decision-window, (darker colour indicates greater voxel z-statistic; All results are reported at Z ≥ 2.57, and cluster-corrected using a resampling procedure resulting in a minimum cluster size of 110 voxels; see **Methods**). **E)** Clusters of voxels with a significant positive relation to the EEG feedback-prediction (3, blue) and bet-prediction (4, yellow) in the feedback-window, with the clusters from the decision-window for comparison. Abbreviations: dSTR, dorsal striatum; GPe, external globus pallidus; IFG, inferior frontal gyrus; INS, insular cortex; pMFC, posterior medial frontal cortex; rlPFC, rostrolateral prefrontal cortex; SPL, superior parietal lobe; vlPFC, ventrolateral prefrontal cortex; vmPFC ventromedial prefrontal cortex; vSTR, ventral striatum.

Applying the same spatial filters estimated within the decision-window onto EEG activity during the feedback-window (i.e. projecting the data through the same “spatial generators” discriminating bet from no-bet responses earlier in the trial), we found the confidence-relevant EEG activity re-emerged. On no-feedback trials, the bet-prediction dissociated bet from no-bet trials (from 0.35 to 0.45 s following feedback, mean t(22) = 2.58, cluster corrected *p* = 0.003, **Figure 2A, right**), suggesting that in the absence of explicit feedback, participants did use their confidence to infer the points they may have gained/lost. On feedback trials, a small window of significant difference between explicit feedback sign was driven primarily by the trials with feedback of −2 (betting on an incorrect response; main effect of feedback sign, mean F(1,22) = 4.96, cluster corrected *p* = 0.035; **Figure 2A, middle**). This could be due to the salience of explicit feedback contradicting confidence, or perhaps the integration of explicit feedback to revise confidence.

To identify the EEG signatures of explicit feedback, the same LDA analysis was applied to discriminate positive (+1 or +2 points) from negative (−1 or −2 points) explicit feedback trials, training within the feedback-window. Spatial filters were first generated for each time-point within the feedback-window, and then a robust individual filter was selected for each participant as the average of five consecutive filters that best discriminated positive from negative feedback trials within the group-level significant time-window (see **Methods, Supplementary Figure 2**). Despite being trained only to discriminate feedback sign, the feedback-prediction showed an interaction between feedback sign and absolute (1 vs. 2) value (from 0.43 to 0.52 s following feedback, mean F(1,22) = 6.81, cluster corrected *p* = 0.006; and for the main effect of feedback sign, the corresponding time-window was from 0.35 to 0.52 s, with mean F(1,22) = 9.62, cluster corrected *p* < 0.001; **Figure 2B, middle**). This reflects a representation of overall outcome value as opposed to just outcome valence. Moreover, the feedback-prediction in the feedback-window predicted response perseveration on the following trial (mean GLM β-weight for repeating stimulus/response interaction = 0.08 ±0.02, t(22) = 8.77, *p* < 0.001), suggesting this analysis was sensitive to the activity relevant for learning from explicit feedback. In addition, the feedback-prediction also dissociated bet from no-bet trials on no-feedback trials (from 0.17 to 0.26 s following feedback, mean F(1,22) = 2.84, cluster corrected *p* = 0.002; **Figure 2B, right**), which could reflect signatures of learning from implicit feedback. Applying the spatial filter back in time onto EEG activity during the decision-window, the feedback prediction also dissociated bet from no-bet trials (from 0.17 to 0.30 s following the response, mean t(22) = 2.96, cluster corrected *p* < 0.001; **Figure 2B, left**). This suggests that some feedback-relevant EEG activity may be present following decisions, either in relation to expected feedback or a direct implementation of early learning updates prior to explicit feedback.

Together these analyses give us four EEG-predictions to inform the GLM analysis of simultaneously acquired fMRI BOLD signal: the bet-prediction that best discriminates bet from no-bet trials in the decision-window (representing post-decision confidence); the bet-prediction that re-emerges at the time of feedback, that discriminates bet from no-bet trials in the absence of explicit feedback (representing implicit outcome value); the feedback-prediction that best discriminates positive from negative feedback in the feedback-window (representing explicit outcome value); and the feedback-prediction in the decision-window, that shows the pattern of EEG activity relevant for discriminating the sign of explicit feedback is present even before explicit feedback is given (representing expected outcome). In addition, regressors on stimulus onset, the bet cue, and the feedback cue were used to capture BOLD related to these externally driven events (**Figure 2C**, see **Methods**). Full details of the results of this GLM can be found in **Supplementary Figures 3-10** and **Supplementary Table 1.** An analysis of the variance inflation factor indicated correlations in these variables were not substantial enough to be problematic for the fMRI analysis.

These EEG-predictions give a fine-grained estimation of the internal variables used to implement behavioural responses, as well as capturing trial-by-trial variability in the neural activity underlying these internal variables. In this way they afford greater explanatory power in capturing meaningful differences between trials with otherwise the same behavioural outcomes. In addition, these predictions disentangle effects in close temporal proximity (within the decision- or feedback-windows), due to the temporal resolution of electrophysiological signals, which would otherwise be difficult to dissociate in sluggish BOLD responses. This greater explanatory power is evidenced through the stronger, and more specific clusters emerging from this analysis compared to an analogous analysis using binary behavioural variables (see **Supplementary Figures 3-10**). Here, we focus on clusters of voxels in which the BOLD related uniquely to the contribution of the EEG-predictions, by leveraging the trial-by-trial variability in the internal representations captured by the LDA analysis of the electrophysiological signals. We expect these EEG-predictions to represent (respectively): 1) post-decision confidence; 2) expected outcome value; 3) explicit outcome value; and 4) implicit outcome value.

In the decision-window, the EEG representation of post-decision confidence (1) was associated with significant clusters in bilateral parietal lobe, posterior medial frontal cortex, inferior frontal gyri, and left rostrolateral prefrontal cortex, reflecting both regions involved in decision-making and the computation of decision confidence, consistent with previous literature [38–41] (**Figure 2D**). In addition, the external globus pallidus was significantly related to post-decision confidence, and these significant voxels largely overlapped with those corresponding to the later representation of explicit outcome value (3). The representation of expected outcome value (2) was, in particular, associated with the parietal operculum, extending into insular cortex, which has previously been associated with the anticipation of reward [42]. In the feedback-window, the representation of explicit outcome value (3) was associated with regions consistent with the valuation network, including the striatum and frontal lobes [43–45], whilst the representation of implicit outcome value (4) was associated with a cluster in left dorsal striatum (**Figure 2E**). A key finding emerging from these results is the presence of a dorsal-ventral spatial gradient within the striatum, where implicit outcome value (based on confidence) is represented more dorsally, while explicit outcome value is represented more ventrally.

Of note, only a subset of these clusters would have arisen without leveraging the trial-wise variability from the EEG-predictions (see **Supplementary Figures 3-10**). In particular, the dorsal striatum cluster corresponding to the re-emergence of confidence as an implicit outcome value estimate in the feedback-window is absent for a comparison of binary bet vs no-bet trials in the feedback window. Similarly, this binary comparison in the decision-window does not capture the relationship between post-decision confidence and the external globus pallidus. Had the endogenous trial-wise variability captured in the EEG-predictions resulted from unrelated noise, these variables would have been less powerful predictors in the fMRI GLM. Rather, the EEG-predictions give a closer approximation of the graded subjective variables underlying behaviour, revealing a richer picture of the BOLD correlates, especially in the basal ganglia.

### Integration of implicit and explicit feedback in external Globus Pallidus

A cluster of voxels centred on the external globus pallidus showed significant relation to both the EEG representation of post-decision confidence and later, in the feedback-window, the EEG representation of explicit outcome value (**Figure 2E**). The external globus pallidus (GPe) receives both cortical and striatal projections as well as sending inhibitory projections to the subthalamic nucleus to control motor (dis)inhibition [46,47], thereby playing an important role in the mediation of motivated behaviour [48]. For this reason, we investigated the post-hoc hypothesis that the external globus pallidus acts as a main subcortical hub for integrating implicit and explicit feedback to drive learning.

We isolated the subcortical voxels that were jointly significant for post-decision confidence and explicit outcome value as the GPe region of interest (ROI) and compared the BOLD response in this ROI with the dorsal striatal cluster related to implicit outcome value, and a ventral striatal cluster most strongly related to explicit outcome value (**Figure 3A**). The connectivity between the striatal ROIs and the GPe was assessed by taking the single-trial Pearson correlation between the voxel-average normalised BOLD over the 10 s following feedback. The average (Fisher transformed) correlation is presented in **Figure 3B** for trials in each feedback condition separately (the BOLD time-courses within the feedback window for each ROI are shown in **Figure 3C**, averaged within each feedback condition, the reader may appreciate the qualitative similarity with the EEG predictions in the feedback-window). Both dorsal and ventral striatum showed strong positive correlation with GPe (dorsal mean = 0.498 ± 0.096; ventral mean = 0.275 ± 0.066; which was significantly less than the dorsal correlation within subjects t(22) = 8.16, *p* < 0.001). For both dorsal and ventral striatum, the correlation with GPe was on average greater the more explicit feedback disagreed with bet-choices (when the participant bet but received negative feedback, compared to when they bet but received positive feedback), although the effect of feedback condition was not significant (comparing positive and negative feedback on bet trials, t(22) = 0.48, *p* = 0.63 for ventral striatum, and t(22) = 1.94, *p* = 0.07 for dorsal stratum). Overall, GPe BOLD covaried with both dorsal and ventral striatum, without substantial difference depending on whether explicit feedback was provided nor on the outcome value.

**Figure 3.**
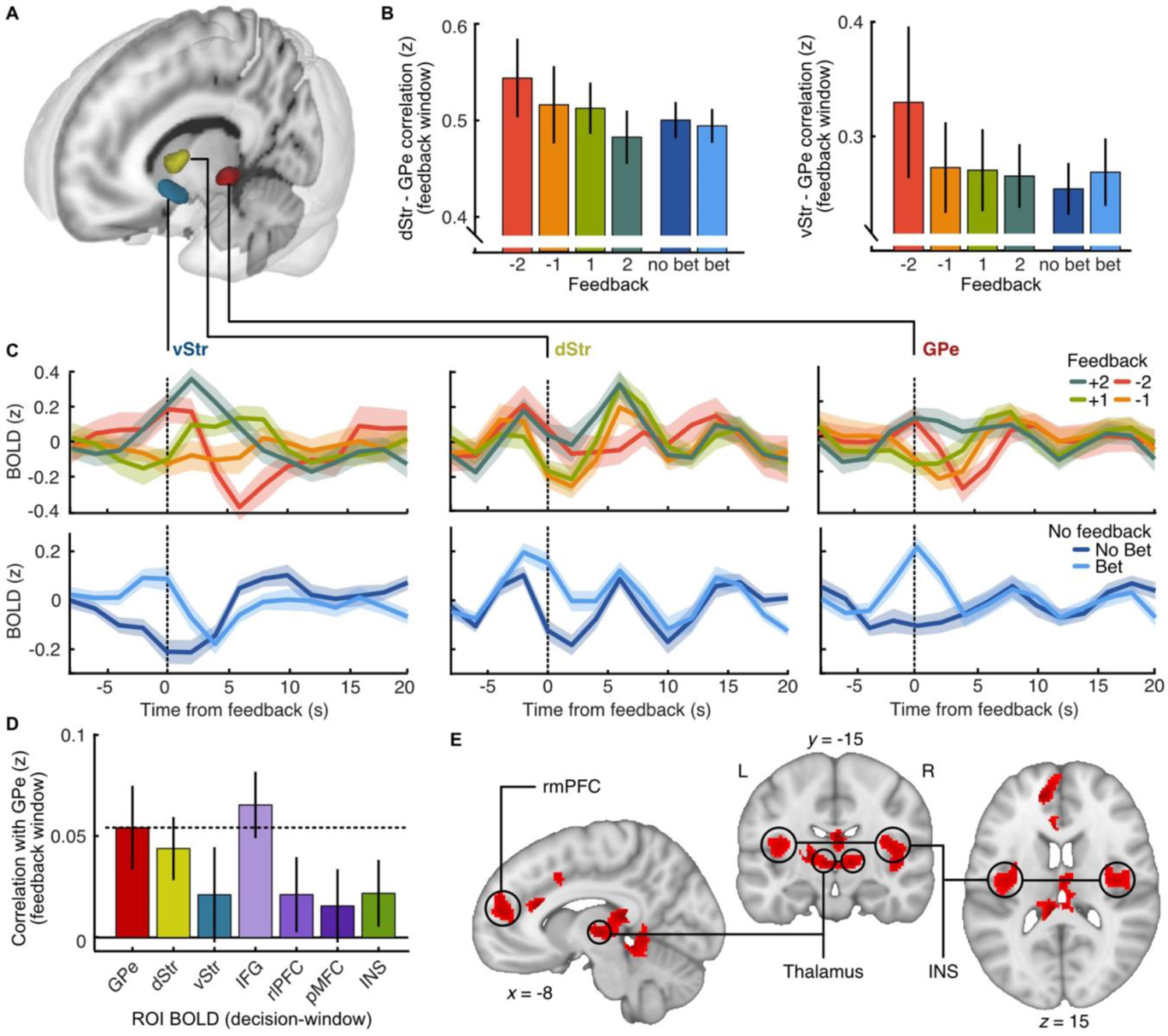
Implicit and explicit feedback signals in the external globus pallidus. **A)** Subcortical regions of interest from the feedback-widow: the dorsal striatum (dStr, yellow, related to the bet-prediction in the feedback-window); the ventral striatum (vStr, blue, related to the feedback-prediction in the feedback-window) and the external globus pallidus (GPe, red, related to both the feedback-prediction in the feedback-window, and the bet-prediction in the decision-window). **B)** Correlation between GPe and dStr (left) and vStr (right) by feedback condition, taken as the Fisher transformed Pearson correlation coefficient over the 10 s following feedback. Error bars show 95% within subject confidence intervals. **C)** Time-course of the normalised BOLD response (averaged over voxels) following feedback by explicit feedback condition (top) and no-feedback condition (bet vs. no-bet, bottom) for vStr (left), dStr (middle), and GPe (right). Shaded regions show 95% within subject confidence intervals. **D)** Correlation between ROI BOLD in the 10 s following the decision with GPe BOLD in the 10 s following feedback. Horizontal dashed line marks the within-region correlation, which can be taken as a benchmark, error bars show 95% within subject confidence intervals. **E)** Significant clusters from the PPI analysis showing the interaction between GPe BOLD following feedback and response perseveration on the following trial. Abbreviations: dSTR, dorsal striatum; GPe, external globus pallidus; IFG, inferior frontal gyrus; INS, insular cortex; pMFC, posterior medial frontal cortex; rlPFC, rostrolateral prefrontal cortex; rmPFC, rostromedial prefrontal cortex; vSTR, ventral striatum.

Next, we assessed how the GPe BOLD in the feedback-window could be driven by earlier BOLD related to post-decision confidence, by taking the correlation in BOLD from up to 10 s following the decision with the later GPe BOLD from up to 10 s following feedback. **Figure 3D** shows the average (Fisher transformed) correlation of the GPe BOLD in the feedback-window with the BOLD in the decision-window for the GPe (itself), dorsal striatum, ventral striatum, three regions related to post-decision confidence (inferior frontal gyrus, left rlPFC, and posterior medial frontal cortex), as well as the insula (related to the expected outcome value). GPe BOLD in the feedback window was significantly correlated with the earlier BOLD in the IFG, and this correlation was on average greater than (though not significantly different from) the earlier BOLD in the GPe itself, suggesting strong evidence for a Granger causal connection. That is, earlier BOLD related to confidence in the IFG continues to drive GPe BOLD responses following feedback. We found no evidence for a difference in this relationship depending on whether explicit feedback was provided (t(22) = 0.912, *p* = 0.37). We formally assessed this relationship within subjects using vector autoregressive models of the BOLD timeseries from the GPe and IFG clusters (see **Methods**). Leave-one-out Granger causal tests showed significant evidence against excluding the lagged IFG BOLD in predicting GPe BOLD in 18/23 participants (median Χ^2^ = 38.3, median *p* = 2.62*e*^−6^, median lag = 12 s, including non-significant participants). Taken together, these results suggest that GPe integrates post-decision cortical confidence and later subcortical outcome value signals, with no substantial difference in this integration depending on whether explicit feedback is provided. This result is consistent with the notion that confidence continues to contribute to the GPe even in the presence of explicit feedback.

Finally, we assessed how the integrated feedback signal encoded in GPe could be used to implement learning, using a Psychophysiological interaction analysis (see **Methods, Supplementary Figure 11**). As the psychological variable we took the interaction between stimulus and response repetition on the following trial, that is, response perseveration. As suggested by behaviour (**Figure 1F)**, we expect positive feedback to be used to increase the likelihood of repeating a response given a stimulus repetition, and negative feedback used to decrease the likelihood of response repetition given stimulus repetition. The psychological variable modelled this interaction, with positive values for switching responses to repeated stimuli following negative feedback (or no-bet responses in the absence of explicit feedback), as well as repeating responses for repeating stimuli following positive feedback (or bet responses in the absence of explicit feedback). The physiological variable was the GPe BOLD from the time of feedback to the start of the next trial. The analysis therefore resolves where increased coupling with GPe (following feedback on trial *n*) results in a stronger influence of (implicit and explicit) feedback on the decision processes on the following trial (pattern of perseveration on trial *n+1*). **Figure 3E** shows the results of this analysis over all trials (both explicit feedback and no-feedback trials), highlighting significant clusters in the thalamus, insula, and the rostromedial prefrontal cortex. The GPe could therefore modulate action value (in medial PFC) and action (dis)inhibition in the thalamus to update decision processes based on both implicit and explicit feedback. Running the PPI analysis separately on explicit feedback trials and no-feedback trials resulted in clusters of significant voxels overlapping with those presented in **Figure 3E**, with no substantial (>2.57) differences in the z-statistics (**Supplementary Figure 9**). This is consistent with the suggestion that the GPe integrates implicit and explicit sources of feedback to drive learning irrespective of the feedback source.

## Discussion

We tested the use of decision confidence for learning in a context where participants could have relied solely on frequent explicit feedback to improve their performance. While at the behavioural level we observed no-feedback trials to be modulated by confidence in a similar manner to explicit feedback trials, at the neural level we found evidence that confidence was integrated to update decision processes even when explicit feedback was provided. These different implicit/explicit sources of value information modulated striatal responses along a dorsal-ventral spatial gradient. We saw evidence that implicit and explicit striatal value signals were integrated in the external globus pallidus, which was significantly modulated by confidence in the decision-window, and explicit outcome value in the feedback-window. Stronger connectivity between the external globus pallidus and the thalamus, insular and frontal cortex predicted the interaction between response perseveration and implicit/explicit feedback sign, emphasising the role of GPe in modulating learning via information flow in the basal ganglia.

Our results highlight the additional processes by which confidence is used for learning. The neural signatures of confidence were not only present following decisions, but re-emerged at the time of feedback. In the decision-window, the confidence prediction estimated from the linear discriminant analysis of the EEG signals corresponded both to brain regions associated with decision-making and the computation of confidence (in line with previous literature [19, 38–41]). Confidence evolved after the decision-window (as has been shown for outcome value signals [44]), such that while the confidence predictions from the feedback-window were estimated using the same spatial generators as the decision-window, the predictions were found to systematically covary with a distinct cluster of BOLD in the dorsal striatum. Importantly, this cluster would not have been identified when contrasting the binary behavioural bet responses in a stand-alone fMRI analysis. The distinction between post-decision confidence and the use of confidence as implicit outcome value was made prominent in this experiment by intermixing explicit- and no-feedback trials with a separate outcome stage, whereas previous experiments examining the neural signatures of confidence in learning have not included an outcome stage [10,11]. We were thus able to delineate separable neural processes associated with the computation of post-decision confidence and the use of confidence as implicit feedback to inform future behaviour.

While the confidence derived implicit feedback signals were isolated to the dorsal striatum, the strongest relation with the EEG explicit outcome value prediction was found in the ventral striatum. This is suggestive of a striatal spatial gradient along the dorsal-ventral axis for implicit vs explicit outcome value, respectively. There are various accounts of the functional division of dorsal and ventral striatum, including associative aspects [33], the type of learning [49], and the learning phase [50,51]. These results add nuances to this discussion: here we see a graded reliance on implicit and explicit feedback representations along the dorsal-ventral axis of the striatum, distinguished across intermixed explicit- and no-feedback trials within the same task. This emphasises the broader distinction of the roles of dorsal and ventral striatum within the context of their cortical inputs [52,53]. Previous studies have shown an integration of post-decision confidence with subjective external value in ventromedial prefrontal cortex [54,55] yet here we suggest separable encoding in the striatum. In this way, the value of an external motivator (whether inferred or explicitly signalled reward) and the value of internal motivations (confidence in performing well at the task) could be encoded separately for the sake of flexible weighting according to the learning context.

Our analysis further suggests these implicit and explicit representations of outcome value are integrated in the external globus pallidus, which corresponded both to post-decision confidence in the decision-window and explicit outcome value in the feedback-window. The external globus pallidus is a central part of the indirect pathway through the basal ganglia [46]. Activation of the indirect pathway was originally thought to increase cortical inhibition via the subthalamic nucleus and thalamus [56]. The functioning of the indirect pathway has since been shown to be more complex [48,57,58], including the modulation of decision-making processes [59,60]. In particular, Lilascharoen and colleagues [61] have shown GPe neurons connecting directly to the parafascicular thalamic nucleus modulating behavioural flexibility. This is in line with our connectivity analysis, where the strength of connectivity between GPe and the thalamus and insular cortex predicted the interaction between response perseveration and implicit/explicit outcome. These findings therefore build on mounting evidence for the cardinal role of GPe in modulating behaviour [47], where in addition, strong projections back from GPe to GABA interneurons in the striatum [57] put the GPe in the position to control information flow throughout the basal ganglia [62].

In summary, these results highlight the ubiquitous use of confidence as implicit feedback for improving future behaviour. The value of implicit feedback is encoded in a distinct manner to that of explicit feedback, where we show a dorsal-ventral spatial gradient within the striatum corresponding to these distinct motivational sources. This illustrates a distinction in striatal coding of value within a single task and learning context. The signals from dorsal and ventral striatum were combined in the external globus pallidus, and facilitated updating of choice behaviour for learning via the thalamus in similar ways following both explicit-feedback and no-feedback trials. This highlights that even when external feedback is available, metacognitive estimates of confidence provide us with nuanced information to update our internal processes for improving behaviour.

## Methods

### Participants

Participants (N = 30; 17 male/ 13 female; age range: 22–33 years) were recruited from the local mailing list and asked to provide written informed consent prior to beginning the experiment. All were right-handed, had normal or corrected to normal vision, and reported no history of neurological problems. The study was approved by the College of Science and Engineering Ethics Committee at the University of Glasgow (CSE01355). Participants were remunerated for their participation in the experiment based on their overall performance (up to a maximum of £10) and an additional fixed payment of £10 for their participation. Due to problems in data collection, two participants were excluded, an additional four participants were excluded for performance below 55% correct, and one participant for too few ‘bet’ responses (betting on fewer than 15% of trials).

### Materials

MRI data was collected using a 3-Tesla Siemens TIM Trio MRI scanner (Siemens, Erlangen, Germany) with a 12-channel head coil. Functional volumes (235 per block) were captured with a T2*-weighted gradient echo, echo-planar imaging sequence (32 interleaved slices, gap: 0.3 mm, voxel size: 3 x 3 x 3 mm, matrix size: 70 x 70, FOV: 210 mm, TE: 30 ms, TR: 2000 ms, flip angle: 80°). An anatomical reference image was acquired using a T1-weighted sequence (192 slices, gap: 0.5 mm, voxel size: 1 x 1 x 1 mm, matrix size: 256 x 256, FOV: 256 mm, TE: 2300 ms, TR: 2.96 ms, flip angle: 9°). For distortion correction, phase and magnitude field maps were acquired (3 x 3 x 3 mm voxels, 32 axial slices, TR: 488 ms, short TE: 4.92 ms, long TE: 7.38 ms).

EEG was collected using a 64-channel (10-20) MR-compatible EEG amplifier system (Brain products, Germany; with Ag/AgCl electrodes, with in-line 10 kOhm surface-mount resistors, EasyCap GmbH, Germany), recorded with Brain Vision software (Brain Vision, USA) at a sampling rate of 5000 Hz. Reference and ground electrodes were built-in between electrodes Fpz and Fz and between electrodes Pz and Oz, respectively. EEG was synchronised with the MRI data acquisition (Syncbox, Brain Products, Germany) with MR triggers stretched to 50 μs using an in-house pulse stretcher. EEG cables were bundled and secured to a cantilever beam running out the back of the bore.

Stimuli were presented on an LCD projector (running at 60 Hz) viewed from the rear of the MR scanner bore via a mirror at a total distance of 95 cm. Stimulus presentation was controlled using Presentation software (Neurobehavioral Systems). Behavioural responses were collected using an MR-compatible button box.

### Task

Each trial was composed of three time-windows, separated by a variable interval of 1-2s: the decision window; the bet window; and the feedback window. Trials were separated by a variable interval of 2-3 s, with a mid-block break of 30 s. Participants performed 6 blocks of 50 trials (300 trials total).

In the decision window, participants performed a variant of the classic random dot kinematogram (RDK) motion direction discrimination task [35]. Participants were presented with an array of 100 white dots placed randomly within a circular aperture (4.8 degrees of visual angle), on a black background. Each frame, dot position was updated according to angular coordinates such that a proportion of dots appeared to move horizontally left or right, while the other dots moved in random directions. The stimulus was presented for approximately 350 ms. The participant was asked to decide whether the dots were moving left or right, and were given up to 1 second to enter their response with a button press. The proportion of dots moving in the coherent direction was first chosen based on a 2-up 1-down staircase procedure prior to the main experiment, and then decreased each block proportionally to the improvement in performance in the previous block (relative increase in proportion correct from the first to the second half of the block), with two exceptions: 1) if proportion correct averaged over the entire block was greater than 0.7, coherence was reduced by ¼ of its current value; or 2) if proportion correct did not increase above 0.5, and was on average less than 0.55, coherence was increased by ½ of its current value).

In the bet window, participants were cued with the text ‘Bet?’ and were given up to 1 second to press a button to bet that their response was correct, doubling the points gained for a correct decision (but also doubling the points lost for an incorrect decision). The absence of a button press within this time was taken as a ‘no-bet’ response.

In the feedback window participants were cued about what type of feedback they would receive. A cyan cue signalled they would receive explicit feedback (text showing +1 point for correct, −1 for incorrect, or +/- 2 in case the participant bet). A magenta cue meant no explicit feedback would be provided, instead participants were shown a question mark cueing them to assess how many points they think they earnt (where points were awarded in the same manner as explicit feedback trials).

If participants failed to respond within 1 second of stimulus offset in the decision window, they were shown the text ‘Time out!’ after a 4-6 second interval (the duration of the bet and feedback windows) and lost 1 point before starting the next trial.

### Pre-processing EEG

EEG preprocessing was performed using EEGLAB [63] and custom scripts [37,63–65] implemented in MATLAB (MathWorks Inc). MR gradient artefacts were removed by subtracting a drifting template (averaged over 80 TRs centred on each TR). Then, a 12 ms median filter was applied, data were downsampled to 1000 Hz, and bandpass filtered between 0.5 and 40 Hz. Blink artefacts were removed by extracting the first principal component from an eye-calibration routine where the participant was instructed to blink, before starting the scanner. Ballistocardiogram (BCG) artefacts were removed by projecting out the principal component closest to a BCG template (created using previous data with the same materials [66]). BCG principal components were extracted from the data low-pass filtered at 4 Hz (the frequency range where BCG artifacts are mainly observed). Since the BCG shares frequency content with the EEG, we adopted this conservative approach to minimise loss of signal power in the underlying EEG signal, where our multivariate discriminant analysis (see below) likely relies on components orthogonal to the BCG artifact. Data were finally re-referenced to the average.

### fMRI

fMRI analyses were conducted using FSL software [67] (FMRIB, fmrib.ox.ac.uk/fsl). The Brain Extraction Tool [68] (BET) was used for brain extraction of the structural images and local field maps. The first 5 volumes of each functional run were discarded. Functional images were slice-time corrected, high-pass filtered at 50 Hz, spatially smoothed (8mm full width half maximum Gaussian kernel), and unwarped using the field maps (using FEAT [69,70]). Motion correction (using MCFLIRT [71]) was performed with parameters saved for later use as nuisance regressors in the GLM analysis. A two-stage registration aligned functional to structural with boundary-based registration, and structural to standard Montreal Neurological Institute (MNI) space with a 12 DOF, nonlinear search.

### Analysis Behaviour

Proportion correct was calculated as the proportion of trials where the observer responded with the true stimulus direction, excluding timed out trials. Statistics were computed on sensitivity (d’; the difference in normalised hit and false-alarm rates). Reaction times were calculated from stimulus offset. Response perseveration was calculated as the probability of repeating a response given a stimulus repeat or stimulus alternation in each feedback condition, to account for different base rates of repetition, this probability was normalised and divided by the overall normalised probability of repeating a response to a repeating/alternating stimulus of each participant, with statistics (2×2 repeated measures ANOVA for the explicit feedback conditions, t-test for bet/no-bet on no-feedback trials) computed on the difference between stimulus repeat and stimulus alternate scores.

### Computational model

To support the behavioural results indicating participants used their confidence to learn, we implemented a simple computational model to describe learning from confidence, in comparison to an equivalent model that does not use learning. Our approach was based on [2] and [12] who suggest that the observer learns by adjusting the weights on the underlying neural responses to maximise reward/accuracy. To simplify, we simulated the effect this weight adjustment would have on the distributions of perceptual evidence underlying decisions [72] (**Supplementary Figure 1**). We assumed that the left and right stimuli resulted in evidence drawn from Gaussian distributions, and that the observer made their decision by comparing this evidence with a criterion. Because the staircase prior to beginning the experiment should have selected stimulus coherence producing ∼75% correct (*d*^′^ ≈ 1), the starting parameters of these distributions were set to unit variance and means at [−0.5, 0.5]. For simplicity, the criterion was fixed at 0. The bet responses were made by comparing the evidence to a secondary criterion, *b* (a free parameter), and the model was fit to minimise the negative log likelihood of participants’ perceptual decisions and bet responses.

The adjustments to the distributions were made in accordance with the reward prediction error on each trial, moderated by the learning rate, α. For example, the mean of the distribution of evidence from a leftward stimulus, μ_L_ on the following trial, *t* + 1, would be adjusted according to the difference between the explicit feedback on that trial, *r*_*t*_, and the expected value, *E*[*V*_*t*_]:

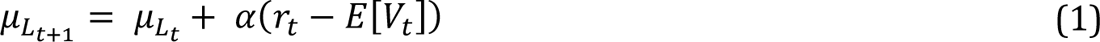

A different learning rate was applied to update the means and standard deviations of the distributions in response to positive and negative reward prediction errors (four free parameters; **Supplementary Figure 1**). For the model using confidence to learn, the expected value on each trial was proportional to the probability of a correct response, based on the cumulative probability density of the distance of the evidence from the criterion, the definition of ideal confidence [73,74]. If the participant bet, the expected value was increased by 1. For the confidence model, the expected value was used as the reward prediction error when no explicit feedback was provided.

This confidence model was compared to an equivalent no-confidence model, which assumed the observer did not use metacognitive information for learning. Thus, the expected value was implemented by means of a Rescorla-Wagner rule [75], starting at 1 (expecting to be correct) and adjusted each trial based on the reward prediction error with a separate learning rate (requiring an additional free parameter). For this model, there was no adjustment made on no-feedback trials. This additional parameter biases the formal model comparison in favour of the simpler, confidence model. Even so, comparison of the model log-likelihood was also in favour of the confidence model, in the majority of participants (16; ∑ ΔNLL = −102.85).

We note that these models could have been improved by implementing additional parameters, for example, starting values for the means of the distributions, or parameterising the perceptual decision criterion (**Supplementary Figure 1**). But the purpose of this modelling exercise was merely to support the behavioural results suggesting participants were using their confidence to learn, as opposed to developing the model frameworks to maximally capture behaviour. Instead, this experiment was designed to focus on the neural mechanisms and leave computational model development to future work.

### EEG

We used a Linear Discriminant Analysis (LDA) to select spatial filters (linear channel weights) that maximally discriminate between bet and no-bet trials, and positive and negative feedback trials (two separate analyses). In each analysis, the LDA was trained on data from a sliding 60 ms window (10 ms intervals) across epochs locked to the time of the response (decision-window) or the time of feedback (feedback-window) using an iterative recursive least squares algorithm for linear logistic regression [76,77]. The sum of the product of the trained weights (*W*, spatial filter) by the multichannel activity (*x*) at trial *t* gives a continuous prediction (*y*) of the binary variable to be discriminated:

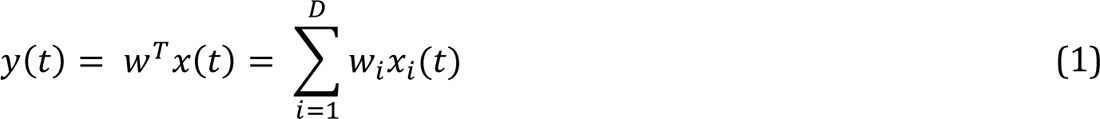

In this way, *y*(*t*) collapses the multichannel data into an aggregate representation that preserves single-trial information while providing a greater signal-to-noise ratio than individual channel activity. Discrimination sensitivity was evaluated using the area under the receiver operating curve (Az) based on predictions (*y*(*t*)) from a leave-one-out cross validation. We generated a robust single spatial filter for each participant, in each of the decision- and feedback-windows, by taking the time-point with the greatest discrimination sensitivity (Az) over a 9-point moving average within the group-level significant time-window (evaluated using a one-sided t-test), and averaging the 5 spatial filters centred on that selected time-point. This procedure is visual depicted in **Supplementary Figure 2**. The spatial filter can be visualised by taking the scalp projections of the discriminating components (the forward model; insets of **Figures 2A** and **B**):

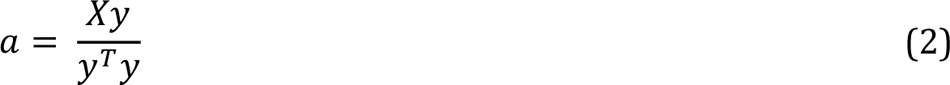

Where *X* is the matrix of channel activity and *y* the vector of predictions. This single spatial filter (for each participant) was then used to generate predictions over time, in both the decision- and feedback-windows. Differences between conditions were tested at each time-point using within-subject t-tests (bet vs no-bet) or ANOVA (feedback 2 sign x 2 value) at an alpha level of 0.05, with cluster correction applied across time [78].

### fMRI

We used an EEG-informed GLM analysis with eight explanatory variables and seven confound variables (the six motion correction parameters, and a custom variable coding for excluded trials and mid-block breaks). Explanatory variables were modelled as boxcar functions convolved with a normalised double-gamma probability density function. The eight explanatory variables were temporally positioned: (1) at stimulus onset, (2) the participant-specific time-point selected for the bet-prediction spatial filter relative to response time, (3) the participant specific time-point where the feedback-prediction best generalised relative to response time, (4 and 5) the time of the bet-cue (for bet and no-bet trials separately), (6) locked to the feedback cue, (7) the participant-specific time-point selected for the feedback-prediction spatial filter relative to feedback time, and (8) the participant specific time-point where the bet-prediction best generalised relative to feedback. All variables were a boxcar function with a duration of 0.1 s, with the exception of the bet cue on bet trials (4; which was extended to the time of the bet-response), and the feedback cue (6; 1 s, the duration of the cue). The boxcar amplitude of the bet-cue on bet trials, the bet-cue on no-bet trials and the feedback-cue variables (4, 5, and 6) was set to 1. The boxcar amplitude of the stimulus onset variable (1) was modulated by the choice reaction time on each trial. The boxcar amplitudes for the remaining four variables were parametrically modulated by the EEG predictions (the bet-prediction in the decision-window, (2); the feedback-prediction in the decision-window, (3); the feedback-prediction in the feedback-window, (7); and the bet-prediction in the feedback-window (8)). These variables correspond to EEG-predictors 1-4 in **Figure 2C**. The EEG feedback-prediction in the feedback-window was only placed on explicit feedback trials, and the bet-prediction in the feedback-window was only placed on no-feedback trials. The design of the eight explanatory variables is visually represented in **Figure 2C**. Note that though the EEG-predictors rely mainly on cortical activity (in close proximity to the sensors) to generate the trial-wise representations, the EEG-informed fMRI analysis can also reveal the contribution of deeper structures that covary systematically with these trial-wise representations. The full results of this analysis are presented in **Supplementary Figures 3-10**, alongside a control analysis using similar variables unmodulated by the EEG-predictors. This control analysis was conducted to be equivalent to the EEG-informed GLM but without the EEG: the bet-prediction in the decision-window was replaced with the binary bet/no-bet response; the feedback-prediction in the decision-window was removed as there is no behavioural equivalent for this; the feedback-prediction in the feedback-window was replaced by the explicit feedback itself (−2/-1/1/2); and the bet-prediction in the feedback-window was replaced again by the binary bet-response. In this way, this analysis reflects what we could have achieved without the EEG, for the interested reader.

Significant clusters were selected using a minimum z-statistic of 2.57 at the group level, and a minimum cluster size of 110 voxels, obtained using a permutation test (described previously [37,64]; the 95^th^ percentile of the empirical null cluster size, calculated over 200 permutations of the EEG feedback-prediction variable (7) order with otherwise the same GLM described above). ROI timeseries, epoched to the closest TR to the time of the decision or feedback, were generated by taking the average z-scored BOLD across ROI voxels after removing values greater than 3.1 standard deviations from the mean over three iterations. Correlations across ROI timeseries were calculated as the average Fisher transformed Pearson correlation across a time-window from 0 to 10 s in the decision-window and the feedback-window (or 0 to 10 s within the feedback-window). The decision- and feedback-windows were separated by 4 to 8 seconds.

We used vector autoregressive models of the full timeseries of GPe and IFG BOLD to test for Granger causality. For each participant, we first selected the appropriate lag in the model based on the Akaike Information Criterion (AIC). The median lag was 6 (12 s) ranging from 2 – 17 (4 – 34 s). All models were found to be stable. Leave-one-out Granger causal tests were conducted within subjects, with 18/23 participants showing significant evidence against the null hypothesis to exclude the history of IFG BOLD in predicting GPe BOLD (over and above the history of GPe BOLD itself). For participants failing to reject the null, the *p*-values ranged from 0.0599 to 0.461, for all other participants, *p* < 0.027 and median *p* = 4.11*e*^−7^.

We also conducted a Psychophysiological Interaction (PPI) analysis [79] to examine how GPe could be used to implement learning. As the physiological variable we took the time-course of the voxel average BOLD in the GPe ROI from the time of feedback to the start of the next trial. The psychological variable described the interaction between stimulus and response repetition that we took as a behavioural signature of learning: trials were coded as +1 for alternating responses to repeating stimuli following negative feedback (or no-bet responses on no-feedback trials), as well as trials with repeating responses to repeating stimuli following positive feedback (or bet responses on no-feedback trials); trials were coded as −1 for repeating a response for a repeated stimulus following negative feedback (or no-bet trials), as well as alternating responses to repeating stimuli following positive feedback (or bet trials). This can be summarised as a positive coding for trials contributing to the positive bars in **Figure 1F** and a negative coding for trials contributing to the negative bars in **Figure 1F**. For further demonstration, an example of the psychological, physiological, and interaction regressors are visually displayed with some annotation in **Supplementary Figure 11**. Clusters of significant voxels were again selected with a statistical threshold of z >= 2.57 and minimum size of 110 voxels. **Figure 3E** shows the results of the analysis across both explicit feedback and no-feedback trials. We also compared the results of the analysis conducted separately on explicit feedback compared to no-feedback trials, taking a difference in z-statistic > 2.57 as evidence for a difference in connectivity. We found no evidence for a difference within the significant regions of the analysis conducted across all trials. These results are presented in full in **Supplementary Figure 12**.

## Acknowledgments

This work was supported in part by the Economic and Social Research Council (ES/L012995/1; MGP) and the European Research Council (865003; MGP). We thank Ralitsa Kostova for assistance with data collection.

## Author contributions

Conceptualisation: MAP, MGP; Methodology: MAP, MGP; Software: TB, MAP, MGP; Formal Analysis: TB; Investigation: MAP; Resources: MGP; Data Curation: TB; Writing – Original Draft: TB; Writing – Review & Editing: TB, MAP, MGP; Visualisation: TB, MGP; Supervision: MGP; Project Administration: MGP; Funding Acquisition: MGP

## Data/code availability

Analysis code, Raw behavioural data, pre-processed EEG, and fMRI Z-statistic maps are available on the Open Science Framework: https://osf.io/q29uf/?view_only=011ffbbc38934678ae85b6a8986e646a

---

**Supplementary Figure 1.**
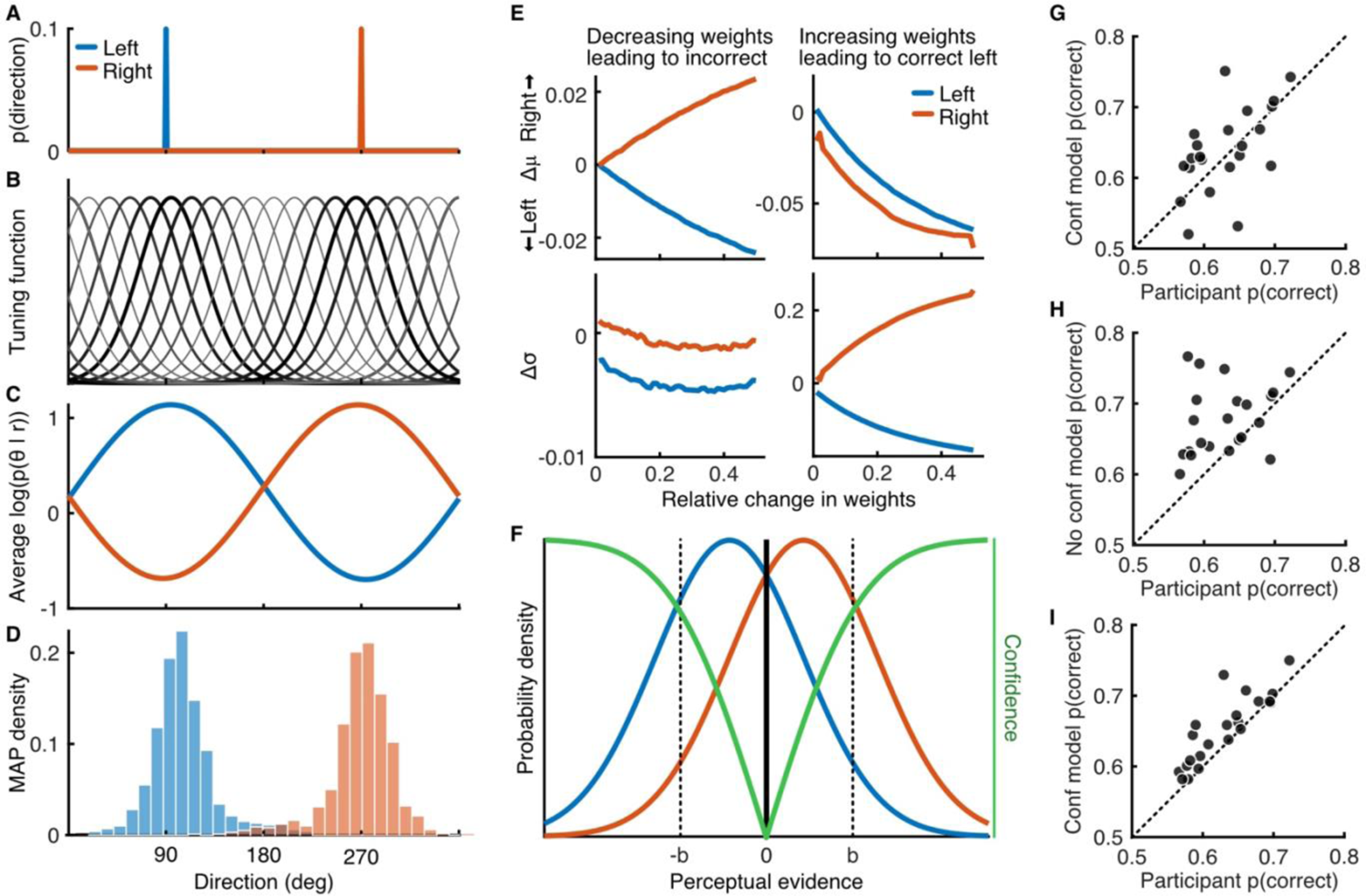
Computational model rationale. The computational model implements reinforcement learning in a perceptual decision-making framework. Previous literature (Law and Gold, 2009) used a reinforcement learning architecture to update weights on neural responses to presented stimuli. We sought to simplify the model by summarising these changes in the weights as changes in the distributions of the resulting perceptual evidence for the response (Dosher and Lu, 1998). We first simulated the responses of MT neurons to the stimuli and the effect of updating the weights. **A)** Probability of dot directions for a left (blue) and right (orange) stimulus with 10% coherent dots. **B)** Simulated population of 20 neurons with circular gaussian (von Mises) tuning functions *f*(θ) with approximate full width half maximum of 70 deg and evenly spaced peaks (Jazayeri & Moveshon, 2006). **C)** Simulating 1000 population responses to each stimulus, where the spike count of each neuron, n, to each dot direction, θ_i_, is drawn from a poission distribution with mean *f*_n_(θ_i_). The log probability of the stimulus, θ_s_, given the population response, r, is the weighted sum of the spike count multiplied by the log of the tuning function (Seung & Sompolinsky, 1993). **D)** The maximum a posteriori (MAP) stimulus direction based on the population response had some variability across simulations, we used this to estimate the mean and variance of the MAP with different weights across neurons. **E)** Following Law and Gold (2009), and Drugowitsch et al., (2019), we assumed the observer should decrease the weights on neurons that drive incorrect responses and increase the weights on neurons that drive correct responses. Those neurons that drive incorrect responses are more likely to be tuned to directions further from the presented stimulus directions (lighter, thinner lines in B), and decreasing the relative weight on these neurons led to greater separation in the mean MAP direction for the two stimuli (top left) and a slight decrease in the variance (bottom left). The neurons contributing most to correct responses have tuning functions centred more closely to each stimulus direction (darker, thicker lines in B). Increasing the weights for neurons contributing to a correct left decision shifts the mean MAP direction leftward (top right) while decreasing the variance in response to a left stimulus and increasing for a rightward stimulus (bottom right). **F)** We implement these changes using the delta rule directly at the level of the perceptual evidence distributions, which we approximate as gaussian for computational ease. The model assumes that on each trial the observer receives a sample of perceptual evidence drawn from the distribution corresponding to the stimulus and makes a left/right decision based on a criterion fixed at 0. The bet response is based on a second criterion at b and -b (free parameter). Confidence (green) estimates the probability of a correct response given the evidence, which is proportional to the cumulative gaussian of the distance of the evidence from the criterion. In the confidence model, this confidence is taken as the expected value, *E*[*V*], on each trial (+1 on a bet trial), and the update is proportional to the reward prediction error, δ = *r* − *E*[*V*], on feedback trials, or the expected value/confidence on no-feedback trials. The learning rate, α, was described by four parameters to apply the updates to the mean and variance of the perceptual evidence distributions in response to positive and negative reward prediction errors separately. **G)** Results of fitting the model to each participants’ choices, showing the predicted proportion correct (across all trials) of the model (y-axis) against the actual proportion correct of the participant (x-axis). **H)** An alternative ‘null hypothesis’ model was formalised for comparison, under this model the participant does not use confidence for learning in any way. On no-feedback trials there is no update. On feedback trials learning proceeds in the same manner as the confidence model, but the expected value is also learnt across trials, starting at 1, and updated according to the reward prediction error, δ at its own learning rate (requiring an additional parameter). This provides a visibly worse fit. **I)** The fit of the confidence model can be improved by adding an additional parameter to account for the fact participants’ initial sensitivity was not always equal to 1 (though the staircase procedure prior to starting experiment aimed to achieve 75% correct, *d*^′^ ≈ 1). This leads to an overall bias for over-performance in the model, which we speculate may be the result of a failure to capture degraded performance due to fatigue in the last block. Potential improvement could also be gained from parameterising other aspects, such as the perceptual decision criterion (fixed at 0), or adding a scaling bias or noise to the underlying confidence estimates. However, the purpose of this exercise was to reaffirm the reader that behaviour can be sufficiently described by a model incorporating confidence for learning (and better than a no-confidence equivalent). We do not intend to develop the computational framework for learning from confidence in this work (we leave this to future work), but rather focus on the neural mechanisms. For this reason, we present the results of the simpler five-parameter model in **G** in the manuscript. Seung, H.S. & Sompolinsky, H. Simple models for reading neuronal population codes. Proc. Natl. Acad. Sci. USA 90, 10749–10753 (1993).

**Supplementary Figure 2.**
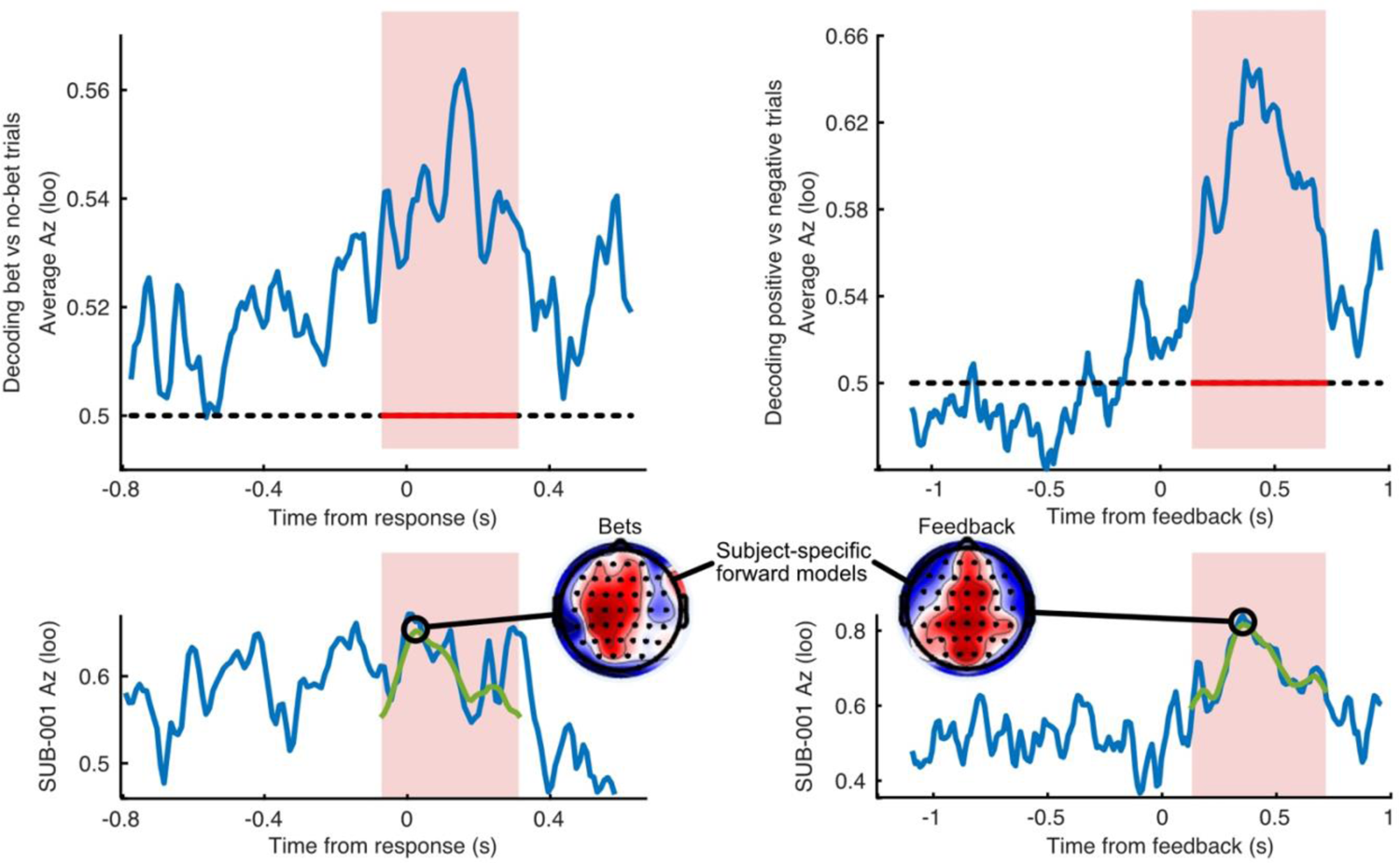
Procedure for isolating subject-specific spatial filters related to representations of confidence and explicit feedback. For each subject, we ran a linear discriminant analysis (LDA) of bet vs no-bet trials in the decision-window and positive vs negative feedback in the feedback window. The analysis was run on data within a sliding window and assessed across time using the area under the ROC (Az) in a leave-one-out (loo) validation procedure. We assessed the group-level significant time window (one-sided t-test) in each case (red shaded areas), and then took the best time-point for individual subjects from a 9-point moving average within this window (green line). The topographies show the resulting forward models (scalp projections of the spatial filter) at this time. The analysis in the Manuscript Figures 2A and 2B is performed by applying these spatial filters over time to generate the predicted value (bet-prediction, or feedback-prediction), for each trial, and then averaging across trials, taking either bet and no-bet trials separately, or explicit feedback trials separately.

**Supplementary Figure 3.**
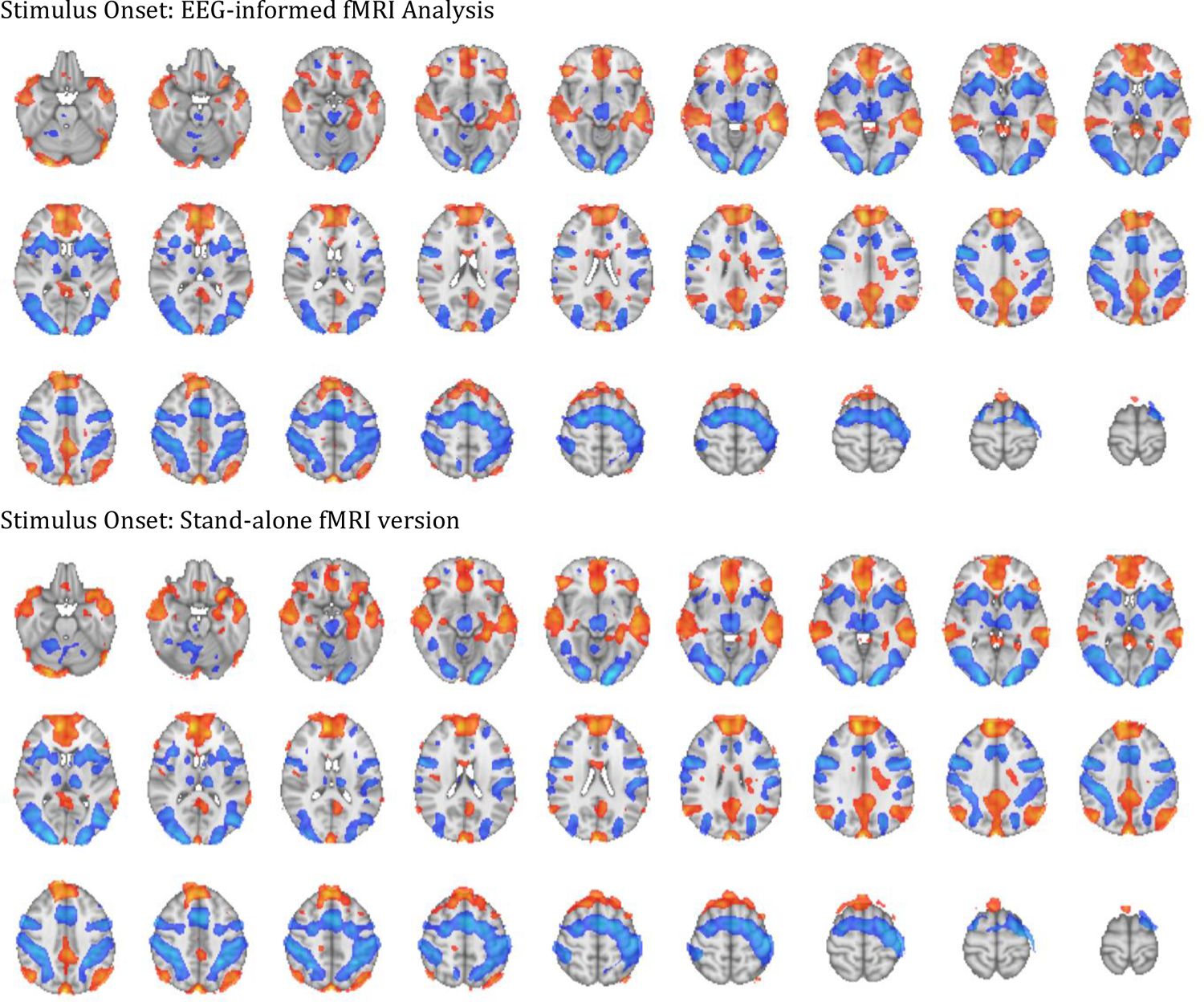
Stimulus onset Z-statistics for the EEG-informed fMRI analysis and the stand-alone fMRI version. The stimulus onset regressor was a boxcar function with duration 0.1 and amplitude modulated by the relative response time of the perceptual decision. Red shows Z-statistics <= −2.57 and Blue, >= 2.57. No minimum cluster size was applied in this display.

**Supplementary Figure 4.**
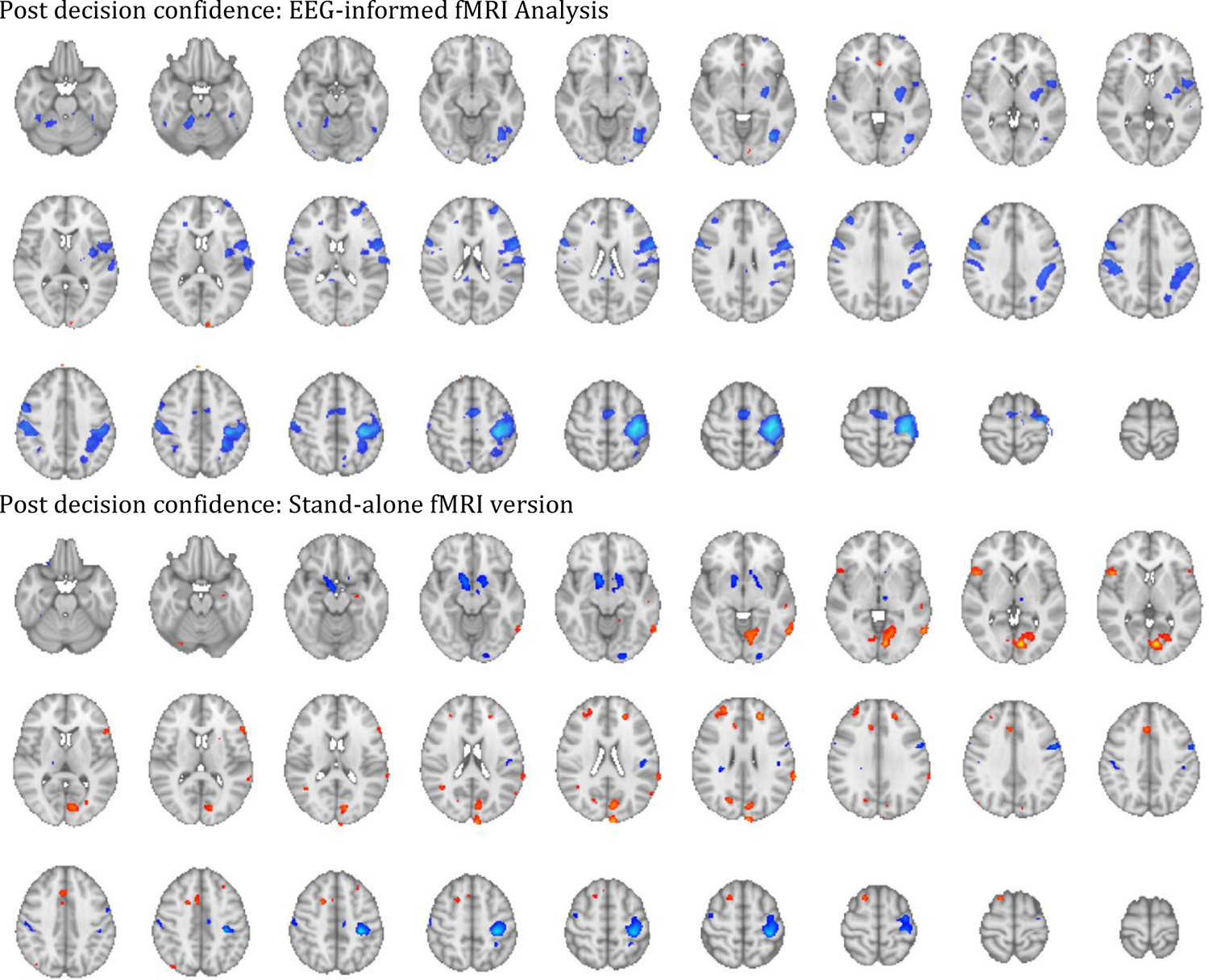
Post-decision confidence Z-statistics for the EEG-informed fMRI analysis (EEG bet-prediction following the perceptual decision) and the stand-alone fMRI version (binary bet = 1, no-bet = −1 following the decision). The regressor was a boxcar function with duration 0.1 and amplitude modulated by the EEG bet-prediction following the decision (or the binary behavioural variable for the stand-alone version). Red shows Z-statistics <= −2.57 and Blue, >= 2.57. No minimum cluster size was applied in this display.

**Supplementary Figure 5.**
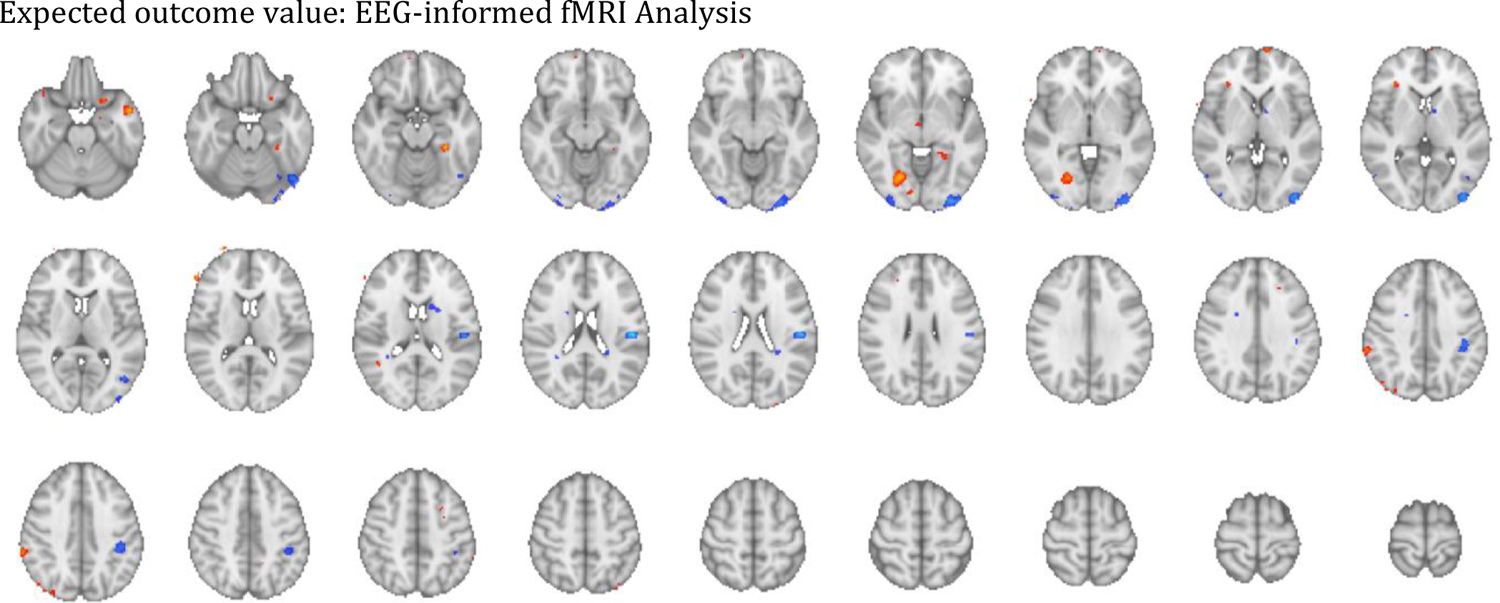
Expected outcome value Z-statistics for the EEG-informed fMRI analysis (EEG feedback-prediction in the decision-window). There is no behavioural proxy for this variable for a stand-alone fMRI analysis. The regressor was a boxcar function with duration 0.1 and amplitude modulated by the EEG feedback-prediction following the decision. Red shows Z-statistics <= −2.57 and Blue, >= 2.57. No minimum cluster size was applied in this display.

**Supplementary Figure 6.**
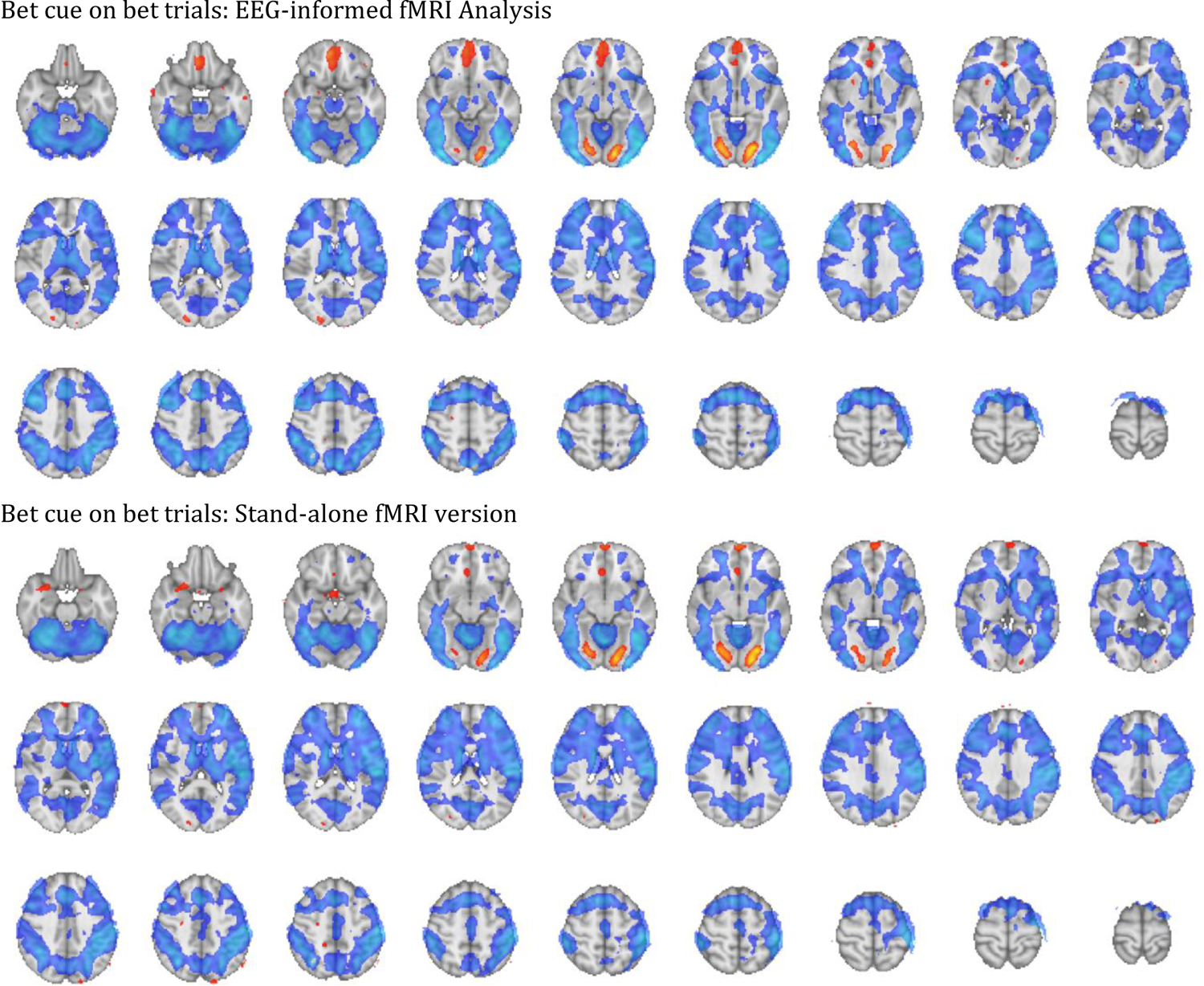
Bet Cue (on bet trials) Z-statistics for the EEG-informed fMRI analysis and the stand-alone fMRI version. The stimulus onset regressor was a boxcar function with duration equal to the bet response time and unmodulated amplitude. Red shows Z-statistics <= −2.57 and Blue, >= 2.57. No minimum cluster size was applied in this display.

**Supplementary Figure 7.**
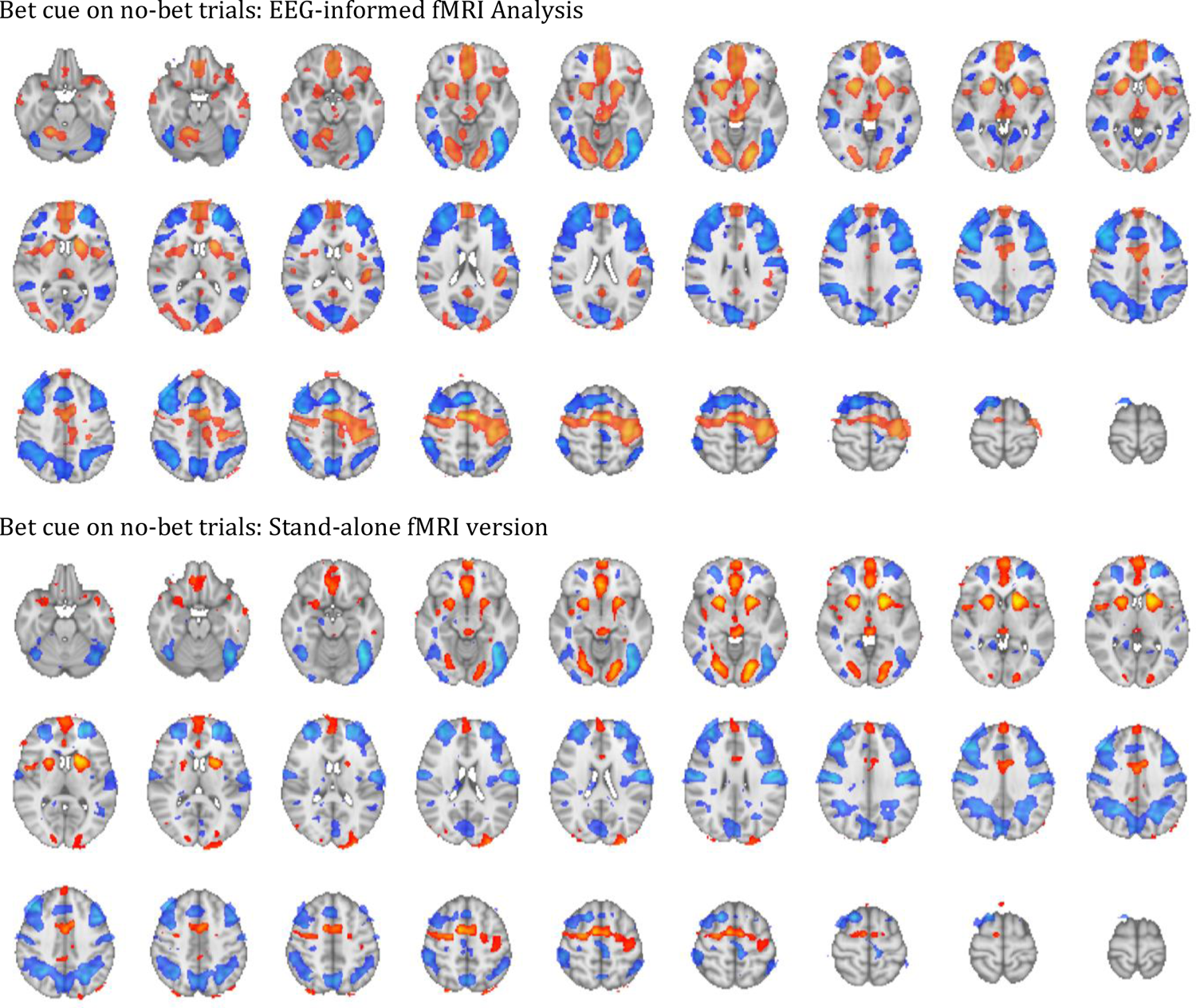
Bet Cue (on no-bet trials) Z-statistics for the EEG-informed fMRI analysis and the stand-alone fMRI version. The stimulus onset regressor was a boxcar function with duration 0.1 and unmodulated amplitude. Red shows Z-statistics <= −2.57 and Blue, >= 2.57. No minimum cluster size was applied in this display.

**Supplementary Figure 8.**
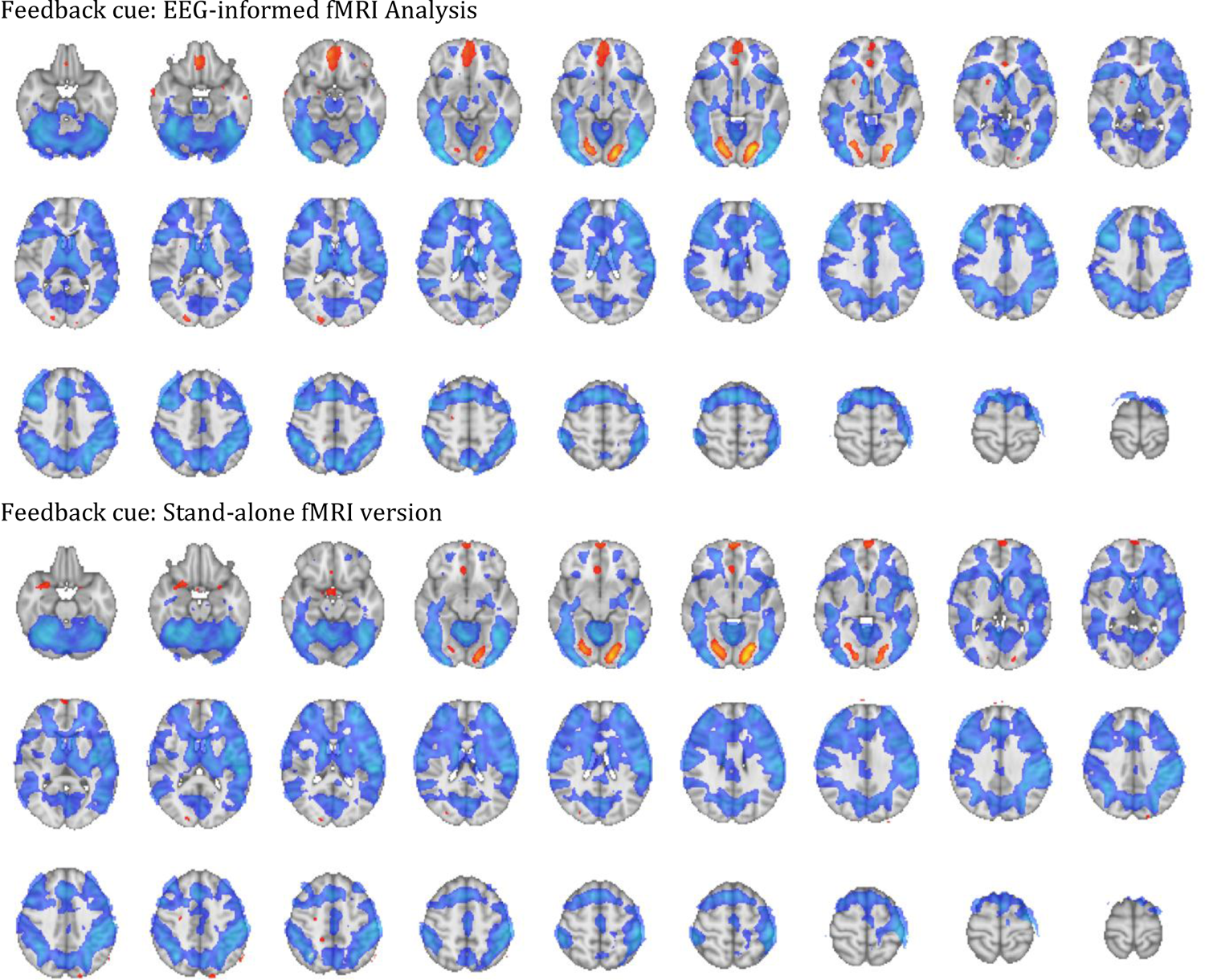
Feedback Cue Z-statistics for the EEG-informed fMRI analysis and the stand-alone fMRI version. The stimulus onset regressor was a boxcar function with duration 0.1 and unmodulated amplitude. Red shows Z-statistics <= −2.57 and Blue, >= 2.57. No minimum cluster size was applied in this display.

**Supplementary Figure 9.**
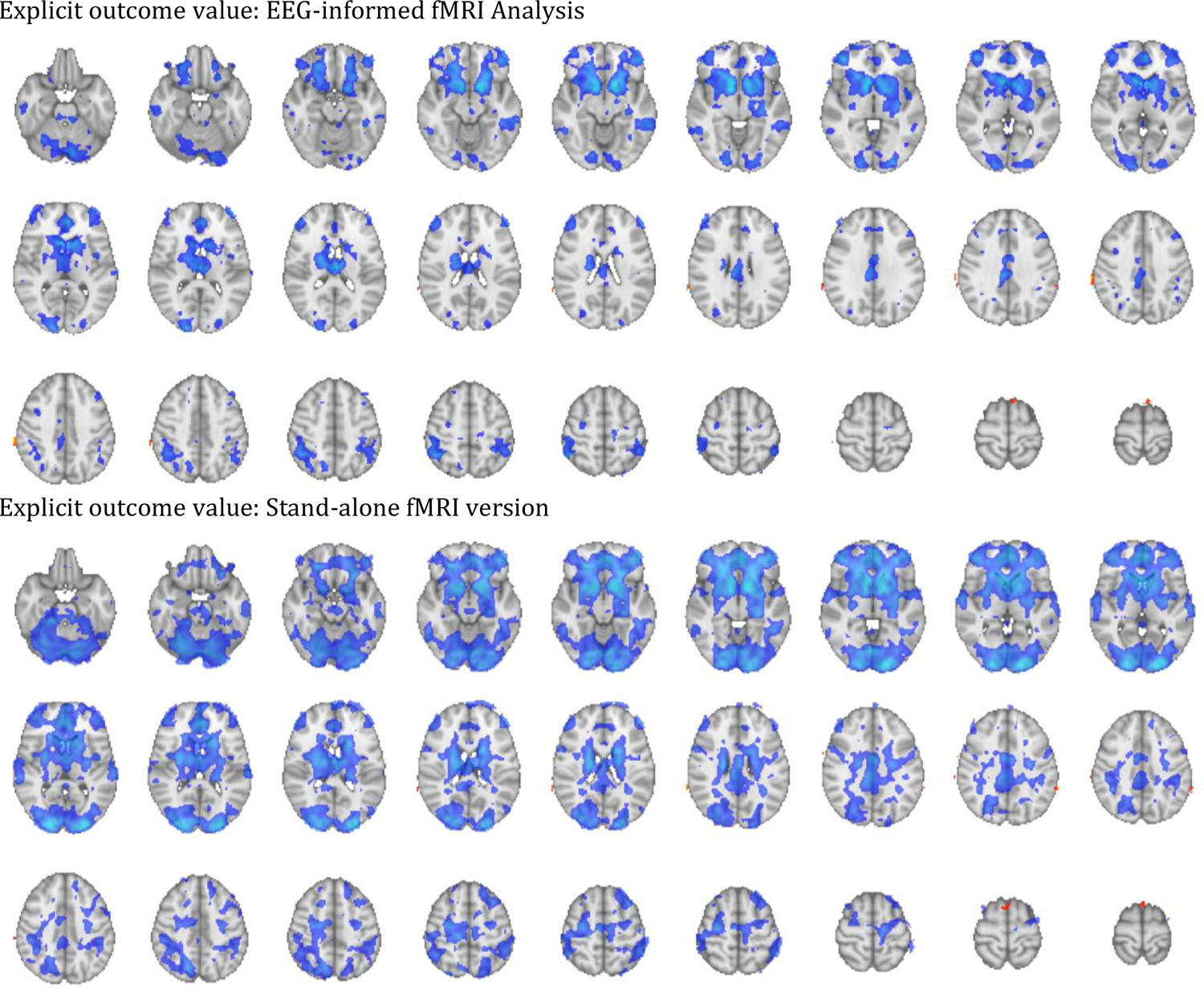
Explicit outcome value Z-statistics for the EEG-informed fMRI analysis (EEG feedback-prediction following explicit feedback) and the stand-alone fMRI version (explicit feedback signed value). The regressor was a boxcar function with duration 0.1 and amplitude modulated by the EEG feedback-prediction following feedback (or the explicit feedback signed value for the stand-alone version). Red shows Z-statistics <= −2.57 and Blue, >= 2.57. No minimum cluster size was applied in this display.

**Supplementary Figure 10.**
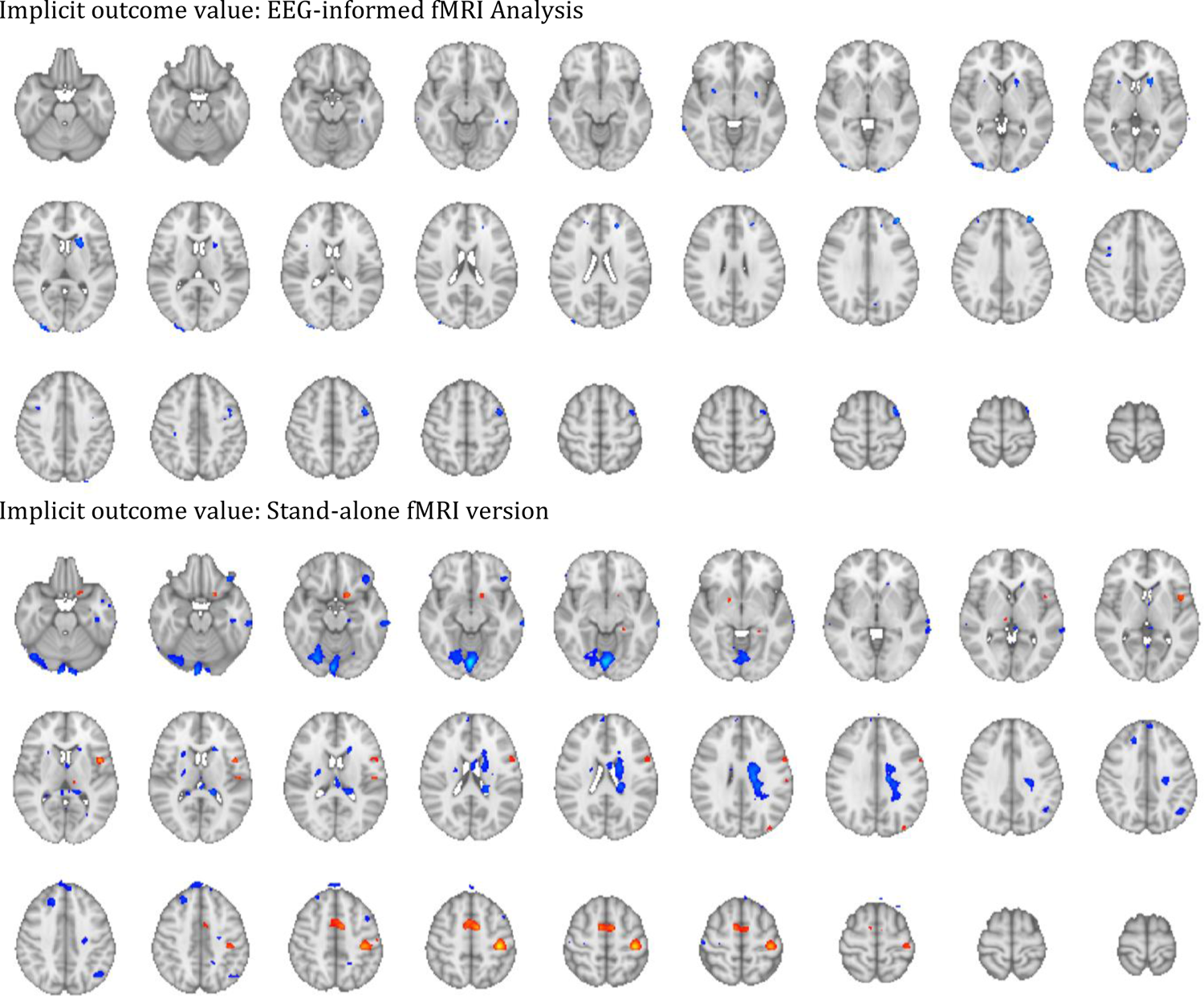
Implicit outcome value Z-statistics for the EEG-informed fMRI analysis (EEG bet-prediction in the feedback-window, on no-feedback trials, top) and the stand-alone fMRI version (binary bet = 1, no-bet = −1 following no-feedback cue, bottom). The regressor was a boxcar function with duration 0.1 and amplitude modulated by the EEG bet-prediction following feedback (or the behavioural variable for the stand-alone version). Red shows Z-statistics <= −2.57 and Blue, >= 2.57. No minimum cluster size was applied in this display.

**Supplementary Figure 11.**
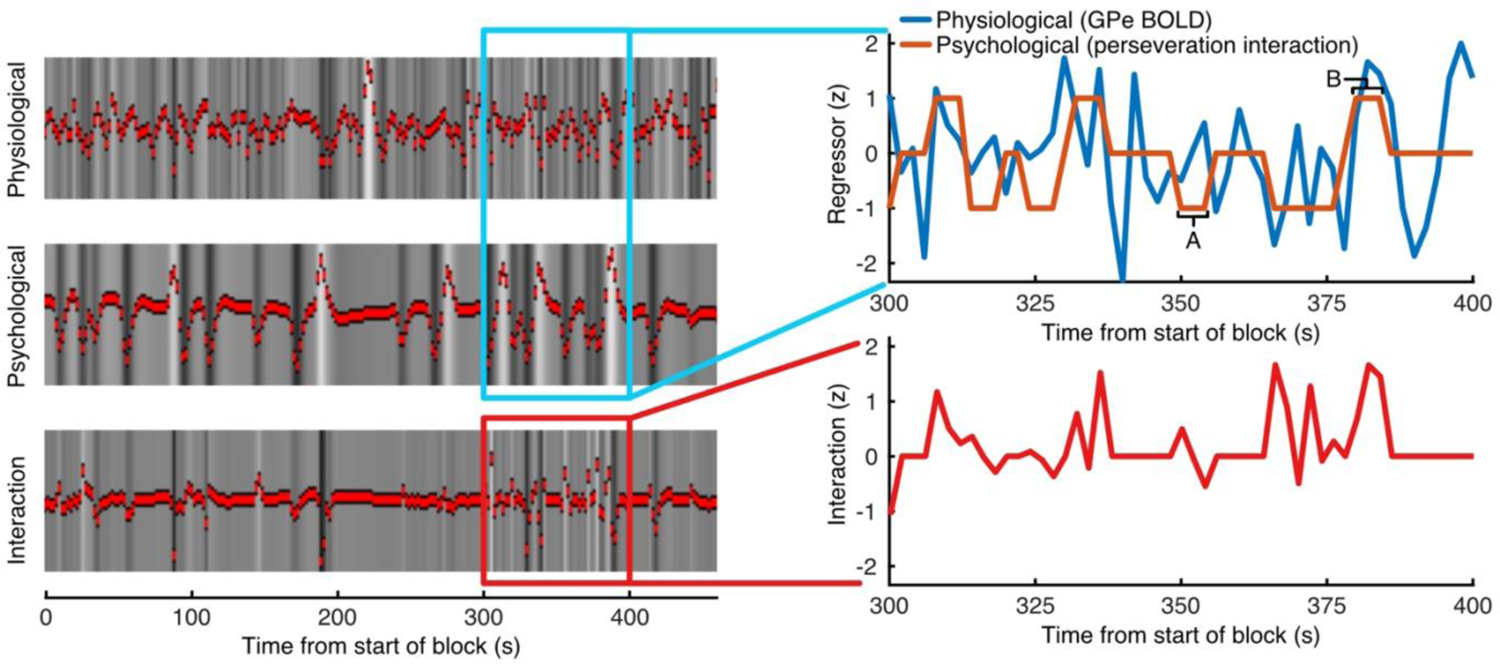
Psychophysiological interaction analysis design. The images on the right show the resulting GLM regressors for one block from one subject. The physiological variable (top) is the GPe BOLD timeseries, taken as the average across voxels in the cluster. The psychological variable (middle) is the perseveration interaction, which is zero except in the time window from feedback over the intertrial interval, where it is set to 1 when the participant behaves according to learning from feedback (alternating responses to repeating stimuli following negative feedback (or no-bet responses on no-feedback trials), as well as trials with repeating responses to repeating stimuli following positive feedback (or bet responses on no-feedback trials), or −1 otherwise (repeating a response for a repeated stimulus following negative feedback (or no-bet trials), as well as alternating responses to repeating stimuli following positive feedback (or bet trials). The interaction is the variable of interest, where clusters of BOLD associated with this variable show increased connectivity with GPe when the participant’s next response will be in line with learning from feedback (or decreased connectivity when the next response is not in line with learning from feedback). The panels to the right zoom in on the variables within the 300-400 s time-window. As a demonstration, the part of the psychological variable marked as A corresponds to the feedback window on trial 41 wherein the participant received negative feedback and on trial 42 the same stimulus direction was repeated and the participant repeated their response. The part of the psychological variable marked as B corresponds to the feedback window on trial 43 wherein the participant received negative feedback and on trial 44 the same stimulus direction was repeated but the participant changed their response.

**Supplementary Figure 12.**
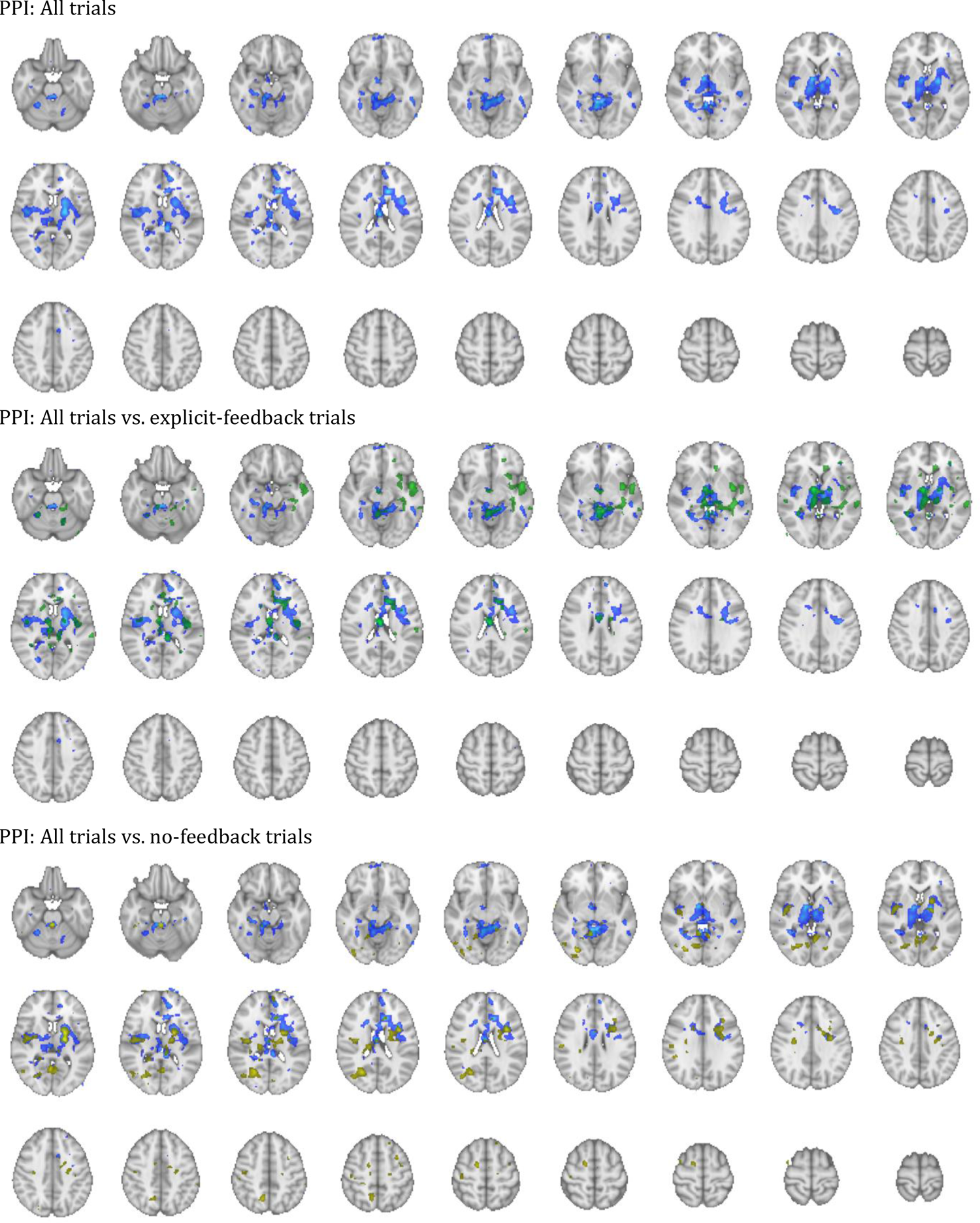
Psychophysiological interaction Z-statistics for the analysis including all trials (top) compared to explicit-feedback trials (middle) and no-feedback trials (bottom). Coloured regions highlight Z >= 2.57 for the interaction between the BOLD time-course from the external globus pallidus region of interest and a variable coding the interaction between response perseveration and feedback (positive for repeating response following positive feedback and alternating response following negative feedback for repeating stimuli, the opposite combination for alternating stimuli, and otherwise negative).

**Supplementary Table 1.**
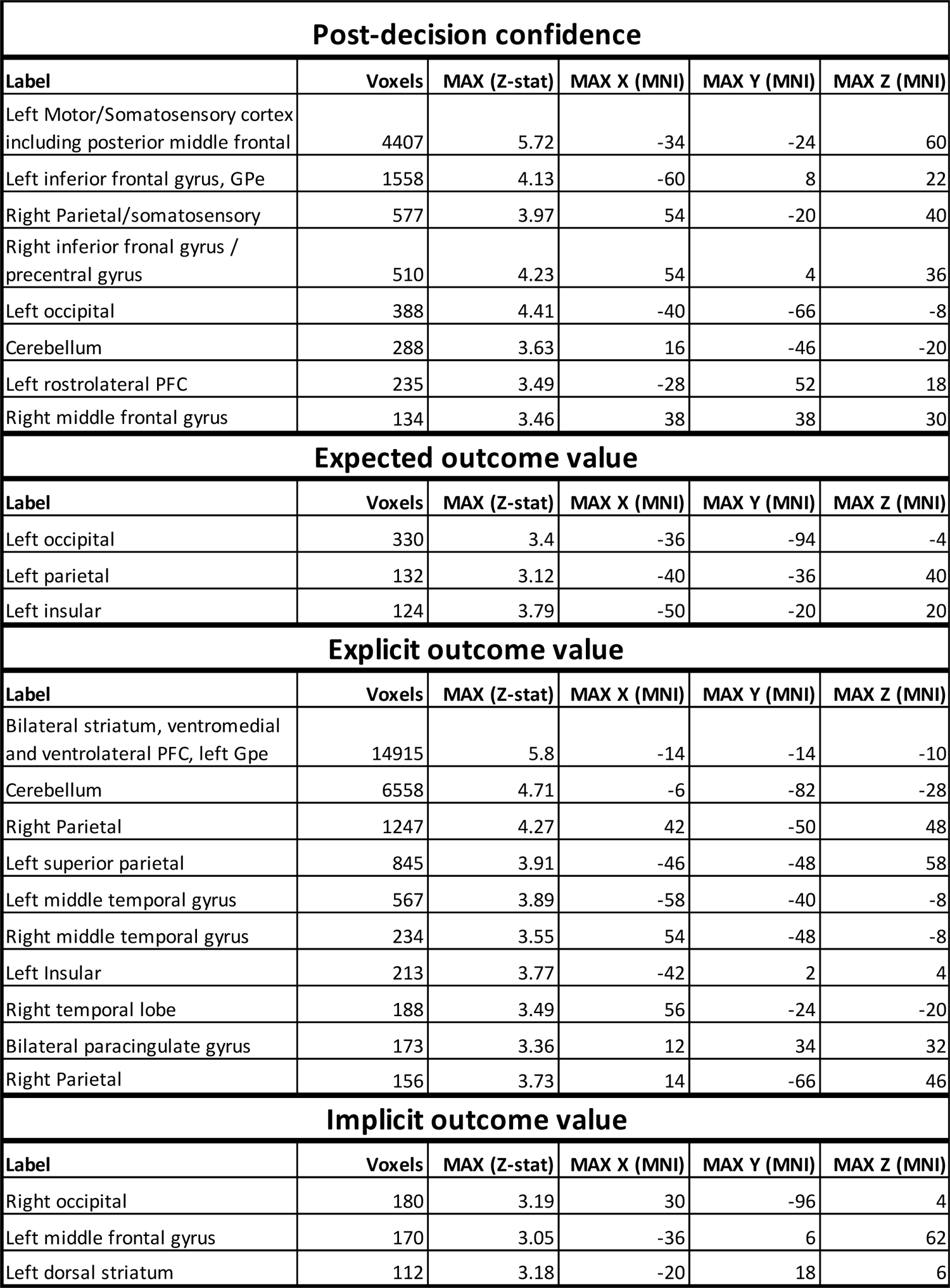
Cluster results for the fMRI GLM analysis (variables of interest only).

## Notes

https://osf.io/q29uf/

### Competing Interest Statement

The authors have declared no competing interest.

