## Supplementary Materials for "Distinct basal ganglia contributions to learning from implicit and explicit value signals in perceptual decision-making"

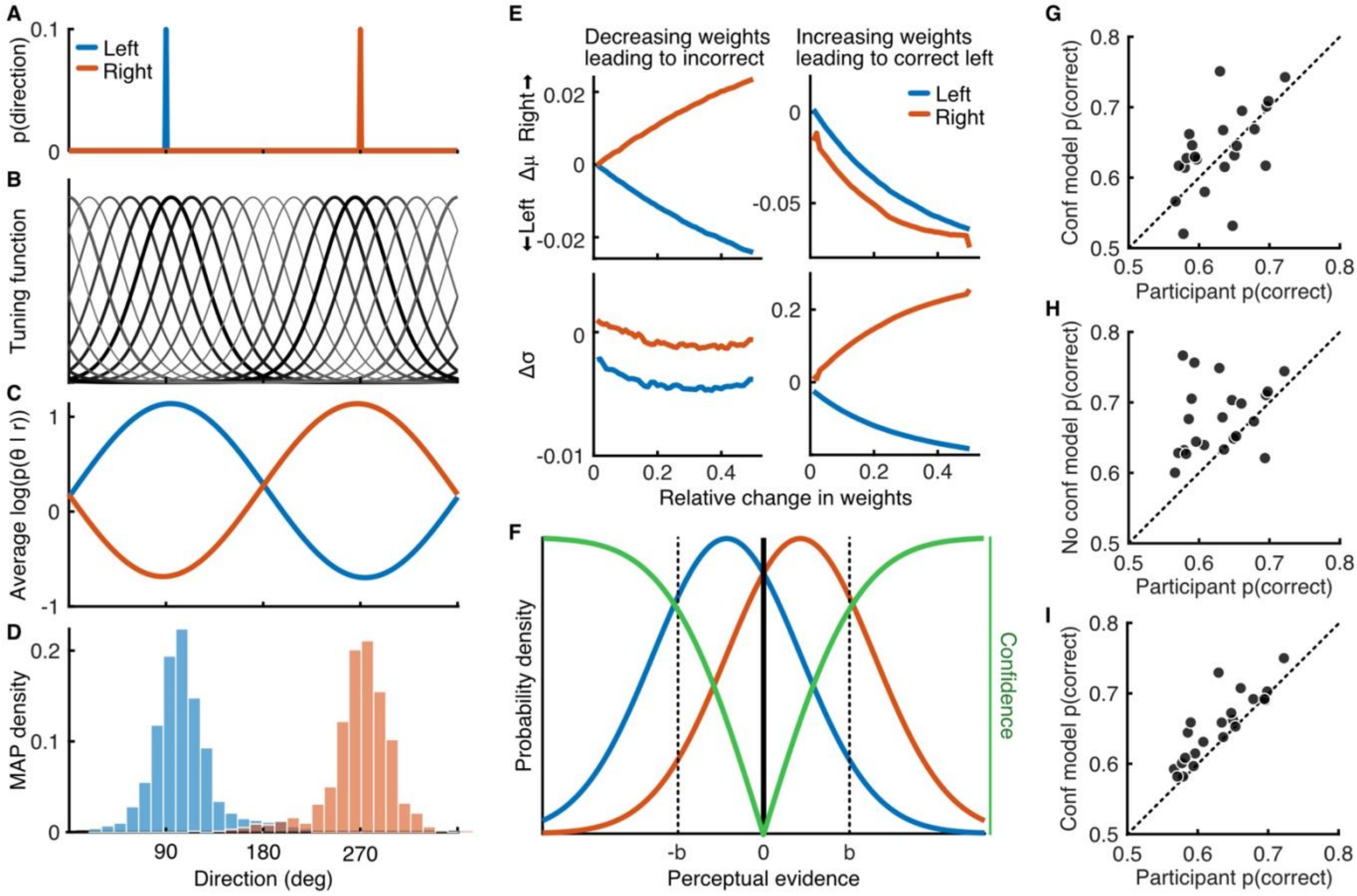


***Supplementary Figure 1. Computational model rationale***. *The computational model implements reinforcement learning in a perceptual decision-making framework. Previous literature (Law and Gold, 2009) used a reinforcement learning architecture to update weights on neural responses to presented stimuli. We sought to simplify the model by summarising these changes in the weights as changes in the distributions of the resulting perceptual evidence for the response (Dosher and Lu, 1998). We first simulated the responses of MT neurons to the stimuli and the effect of updating the weights.* ***A)*** *Probability of dot directions for a left (blue) and right (orange) stimulus with 10% coherent dots.* ***B)*** *Simulated population of 20 neurons with circular gaussian (von Mises) tuning functions $f(\theta)$ with approximate full width half maximum of 70 deg and evenly spaced peaks (Jazayeri & Moveshon, 2006).* ***C)*** *Simulating 1000 population responses to each stimulus, where the spike count of each neuron, n, to each dot direction, $\theta_i$, is drawn from a poission distribution with mean $f_n(\theta_i)$. The log probability of the stimulus, $\theta_s$, given the population response, r, is the weighted sum of the spike count multiplied by the log of the tuning function (Seung & Sompolinsky, 1993).* ***D)*** *The maximum a posteriori (MAP) stimulus direction based on the population response had some variability across simulations, we used this to estimate the mean and variance of the MAP with different weights across neurons.* ***E)*** *Following Law and Gold (2009), and Drugowitsch et al., (2019), we assumed the observer should decrease the weights on neurons that drive incorrect responses and increase the weights on neurons that drive correct responses. Those neurons that drive incorrect responses are more likely to be tuned to directions further from the presented stimulus directions (lighter, thinner lines in B), and decreasing the relative weight on these neurons led to greater separation in the mean MAP direction for the two stimuli (top left) and a slight decrease in the variance (bottom left). The neurons contributing most to correct responses have tuning functions centred more closely to each stimulus direction (darker, thicker lines in B). Increasing the weights for neurons contributing to a correct left decision shifts the mean MAP direction leftward (top right) while decreasing the variance in response to a left stimulus and increasing for a rightward stimulus (bottom right).* ***F)*** *We implement these changes using the delta rule directly at the level of the perceptual evidence distributions, which we approximate as gaussian for computational ease. The model assumes that on each trial the observer receives a sample of perceptual evidence drawn from the distribution corresponding to the stimulus and makes a left/right decision based on a criterion fixed at 0. The bet response is based on a second criterion at b and -b (free parameter). Confidence (green) estimates the probability of a correct response given the evidence, which is proportional to the cumulative gaussian of the distance of the evidence from the criterion. In the confidence model, this confidence is taken as the expected value, $E[V]$, on each trial (+1 on a bet trial), and the update is proportional to the reward prediction error, $\delta = r - E[V]$, on feedback trials, or the expected value/confidence on no-feedback trials. The learning rate, $\alpha$, was described by four parameters to apply the updates to the mean and variance of*

Seung, H.S. & Sompolinsky, H. Simple models for reading neuronal population codes. Proc. Natl. Acad. Sci. USA 90, 10749–10753 (1993).

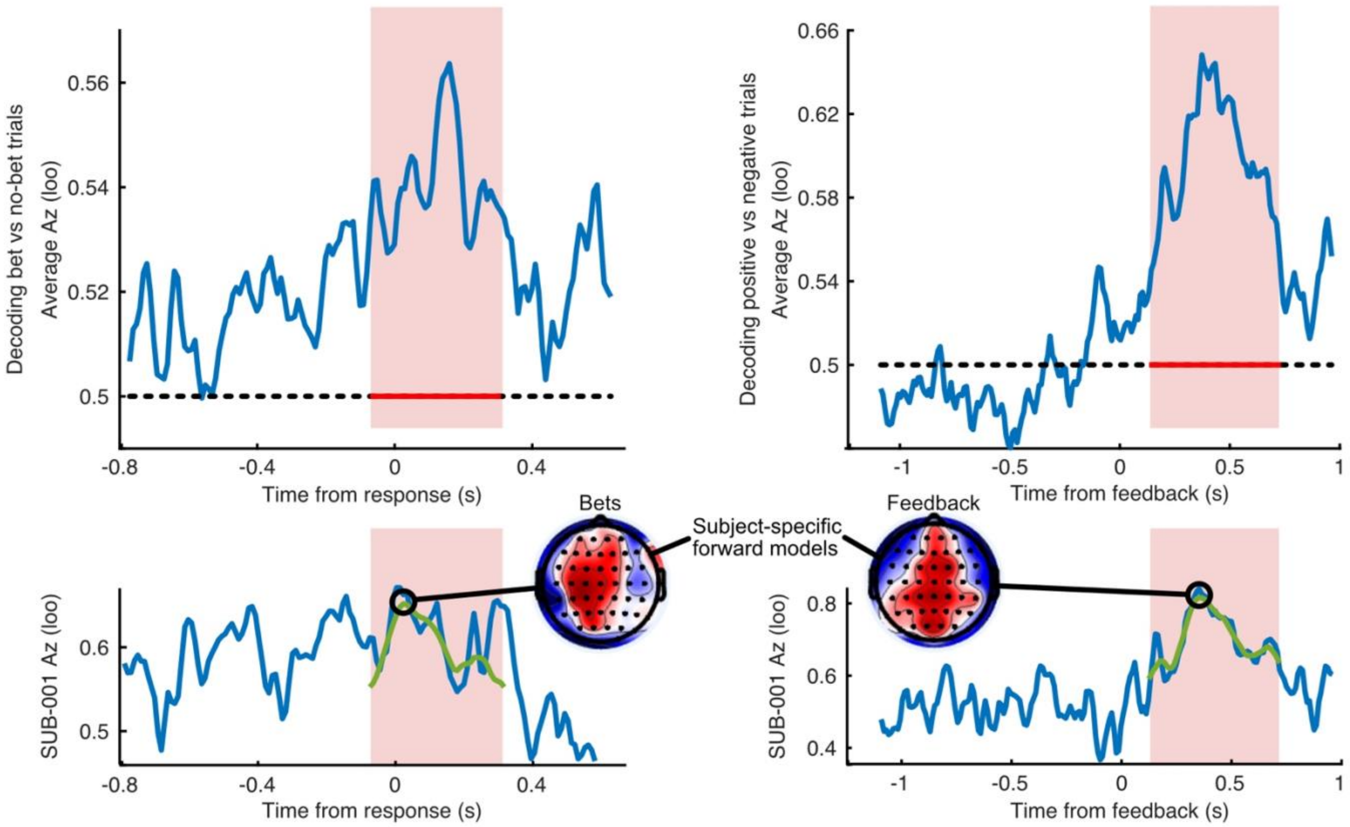


***Supplementary Figure 2.*** *Procedure for isolating subject-specific spatial filters related to representations of confidence and explicit feedback. For each subject, we ran a linear discriminant analysis (LDA) of bet vs no-bet trials in the decision-window and positive vs negative feedback in the feedback window. The analysis was run on data within a sliding window and assessed across time using the area under the ROC (Az) in a leave-one-out (loo) validation procedure. We assessed the group-level significant time window (one-sided t-test) in each case (red shaded areas), and then took the best time-point for individual subjects from a 9-point moving average within this window (green line). The topographies show the resulting forward models (scalp projections of the spatial filter) at this time. The analysis in the Manuscript Figures 2A and 2B is performed by applying these spatial filters over time to generate the predicted value (bet-prediction, or feedback-prediction), for each trial, and then averaging across trials, taking either bet and no-bet trials separately, or explicit feedback trials separately.*

Stimulus Onset: EEG-informed fMRI Analysis

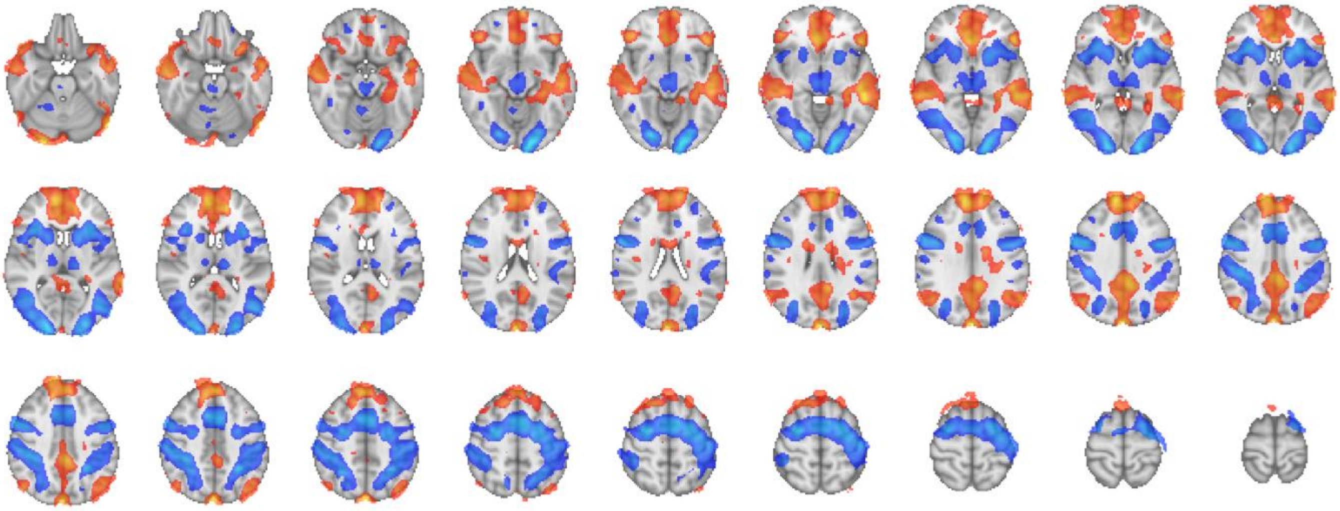

Stimulus Onset: Stand-alone fMRI version

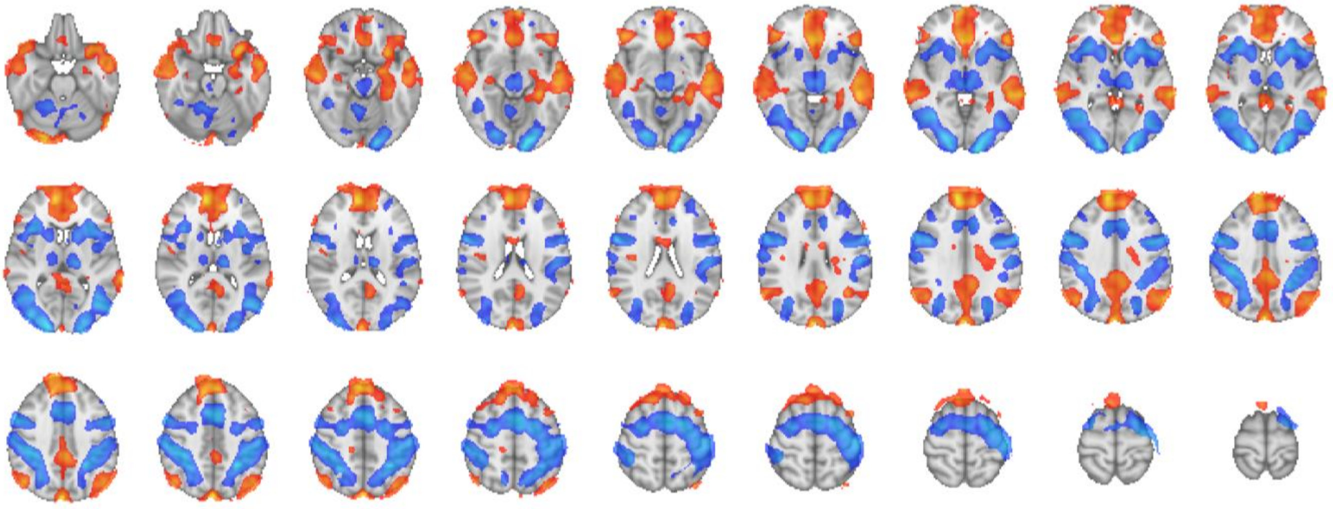

***Supplementary Figure 3.*** *Stimulus onset Z-statistics for the EEG-informed fMRI analysis and the stand-alone fMRI version. The stimulus onset regressor was a boxcar function with duration 0.1 and amplitude modulated by the relative response time of the perceptual decision. Red shows Z-statistics <= -2.57 and Blue, >= 2.57. No minimum cluster size was applied in this display.*

Post decision confidence: EEG-informed fMRI Analysis

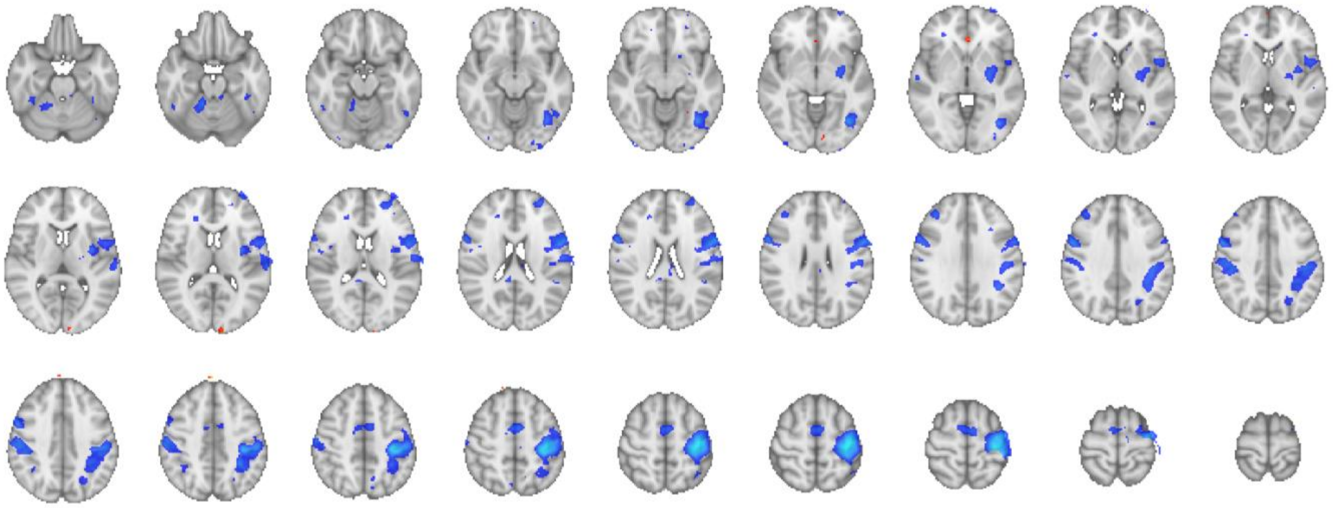

Post decision confidence: Stand-alone fMRI version

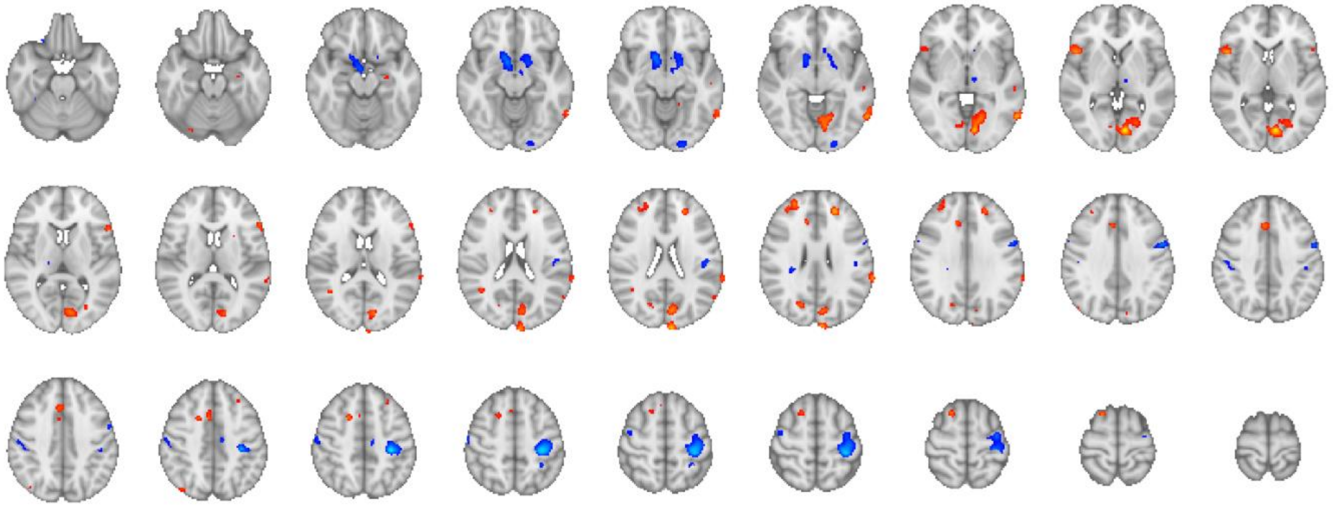

***Supplementary Figure 4.*** *Post-decision confidence Z-statistics for the EEG-informed fMRI analysis (EEG bet-prediction following the perceptual decision) and the stand-alone fMRI version (binary bet = 1, no-bet = -1 following the decision). The regressor was a boxcar function with duration 0.1 and amplitude modulated by the EEG bet-prediction following the decision (or the binary behavioural variable for the stand-alone version). Red shows Z-statistics <= -2.57 and Blue, >= 2.57. No minimum cluster size was applied in this display.*

Expected outcome value: EEG-informed fMRI Analysis

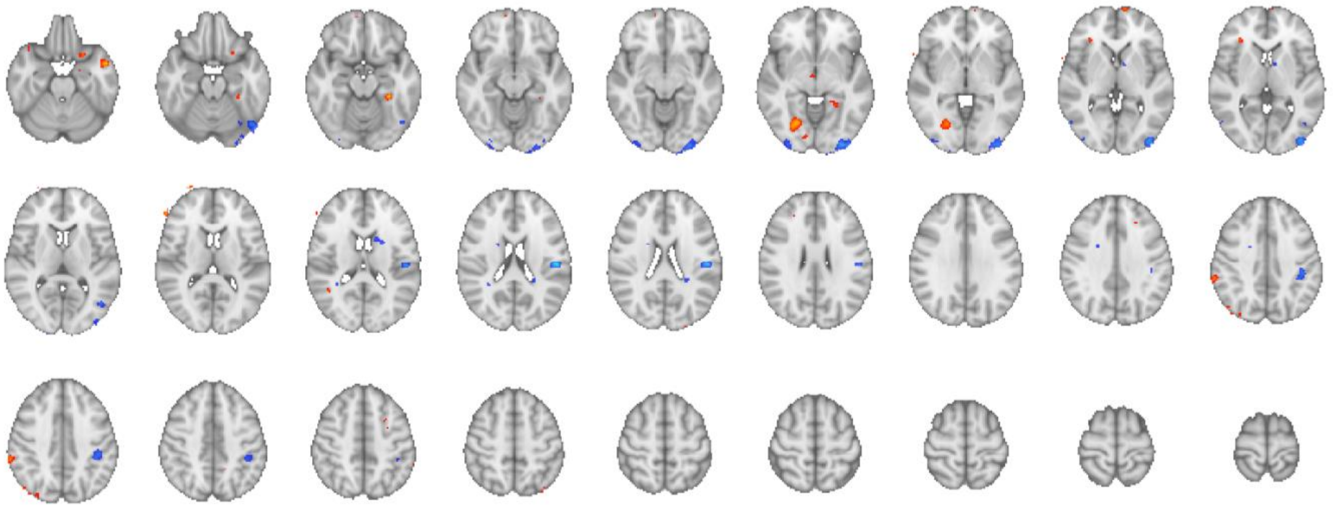

***Supplementary Figure 5.*** *Expected outcome value Z-statistics for the EEG-informed fMRI analysis (EEG feedback-prediction in the decision-window). There is no behavioural proxy for this variable for a stand-alone fMRI analysis. The regressor was a boxcar function with duration 0.1 and amplitude modulated by the EEG feedback-prediction following the decision. Red shows Z-statistics <= -2.57 and Blue, >= 2.57. No minimum cluster size was applied in this display.*

Bet cue on bet trials: EEG-informed fMRI Analysis

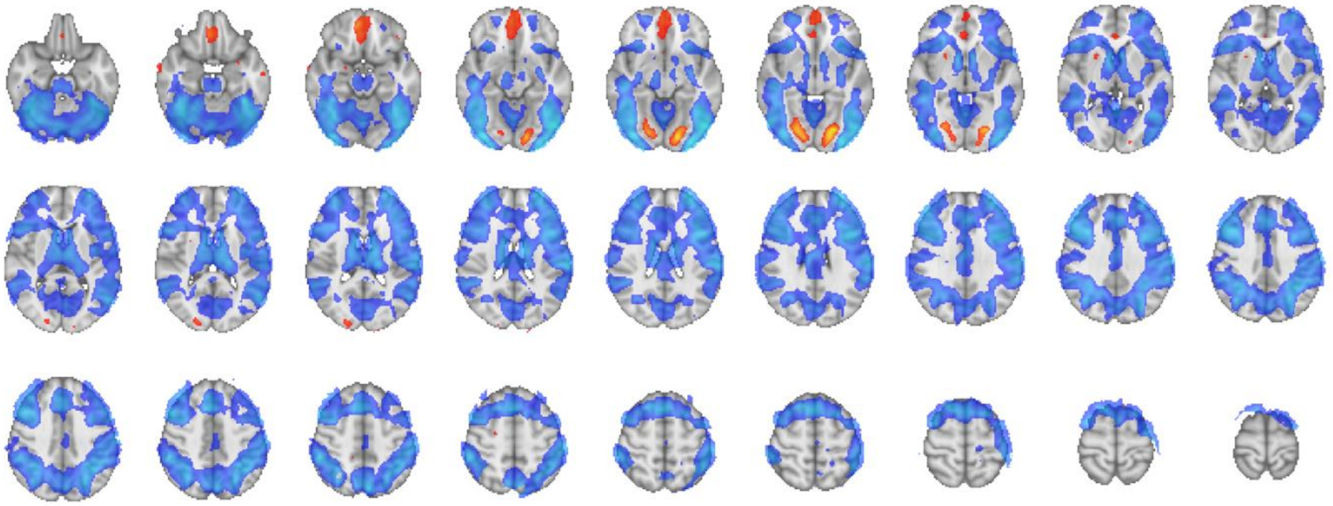

Bet cue on bet trials: Stand-alone fMRI version

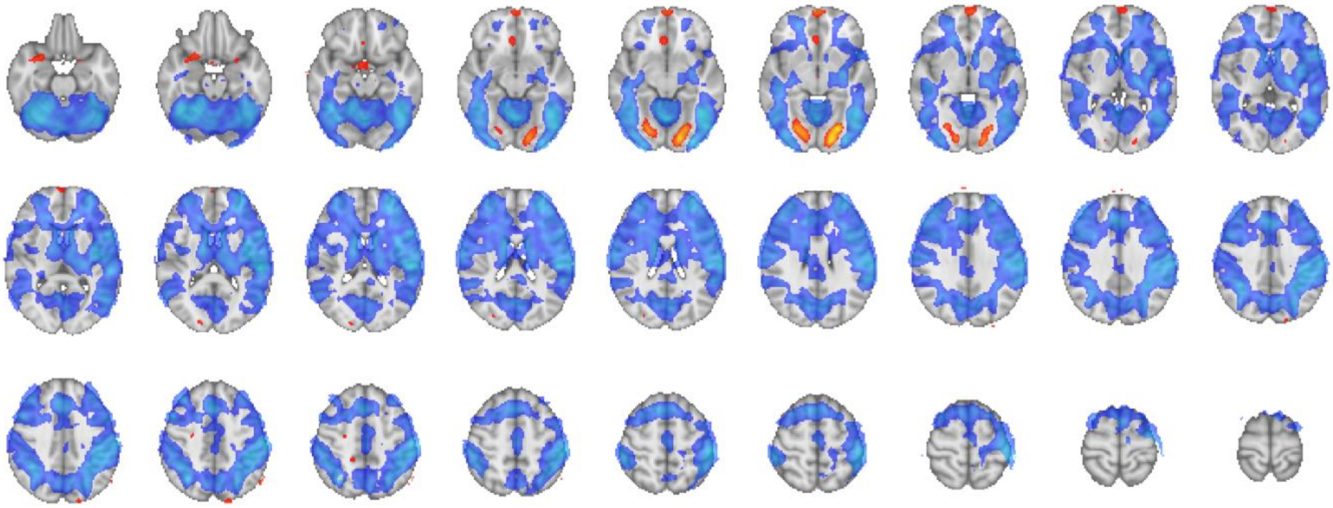

***Supplementary Figure 6.*** *Bet Cue (on bet trials) Z-statistics for the EEG-informed fMRI analysis and the stand-alone fMRI version. The stimulus onset regressor was a boxcar function with duration equal to the bet response time and unmodulated amplitude. Red shows Z-statistics <= -2.57 and Blue, >= 2.57. No minimum cluster size was applied in this display.*

Bet cue on no-bet trials: EEG-informed fMRI Analysis

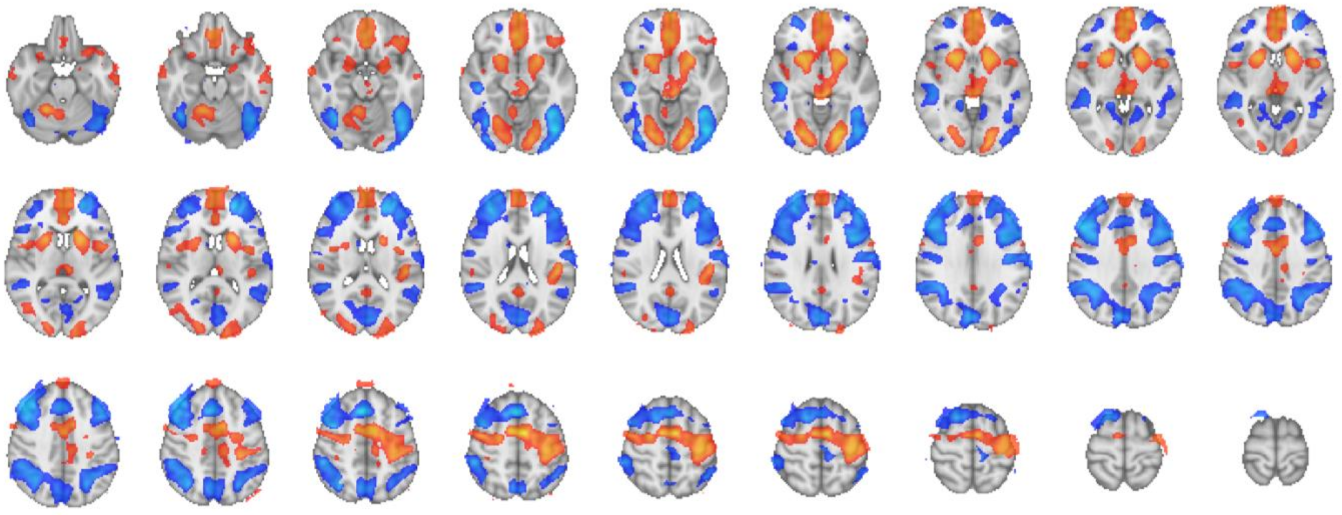

Bet cue on no-bet trials: Stand-alone fMRI version

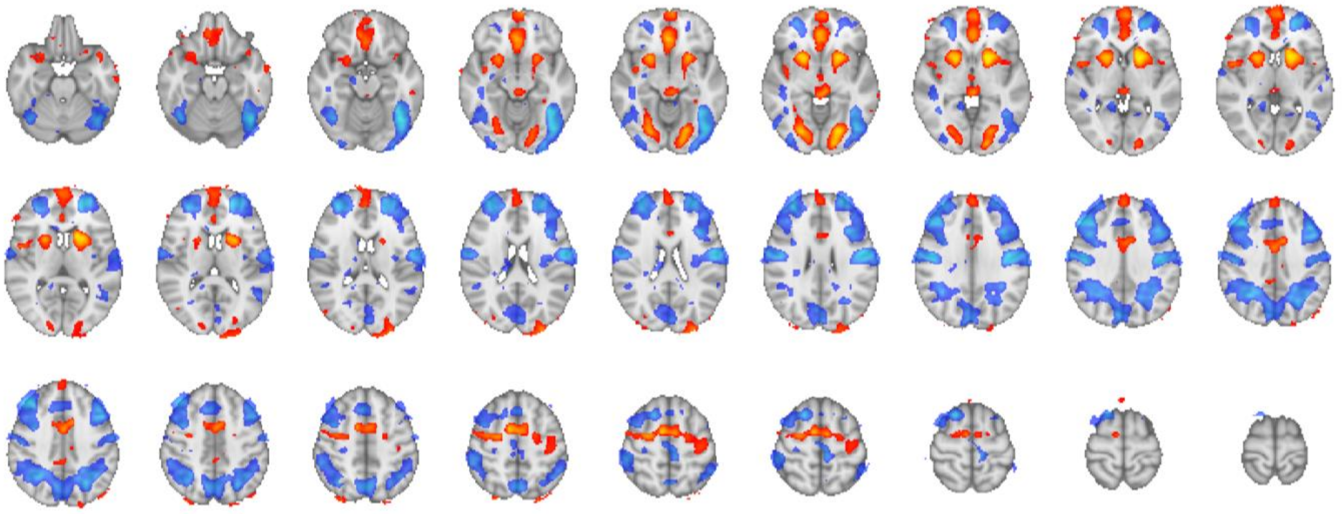

***Supplementary Figure 7.*** *Bet Cue (on no-bet trials) Z-statistics for the EEG-informed fMRI analysis and the stand-alone fMRI version. The stimulus onset regressor was a boxcar function with duration 0.1 and unmodulated amplitude. Red shows Z-statistics <= -2.57 and Blue, >= 2.57. No minimum cluster size was applied in this display.*

Feedback cue: EEG-informed fMRI Analysis

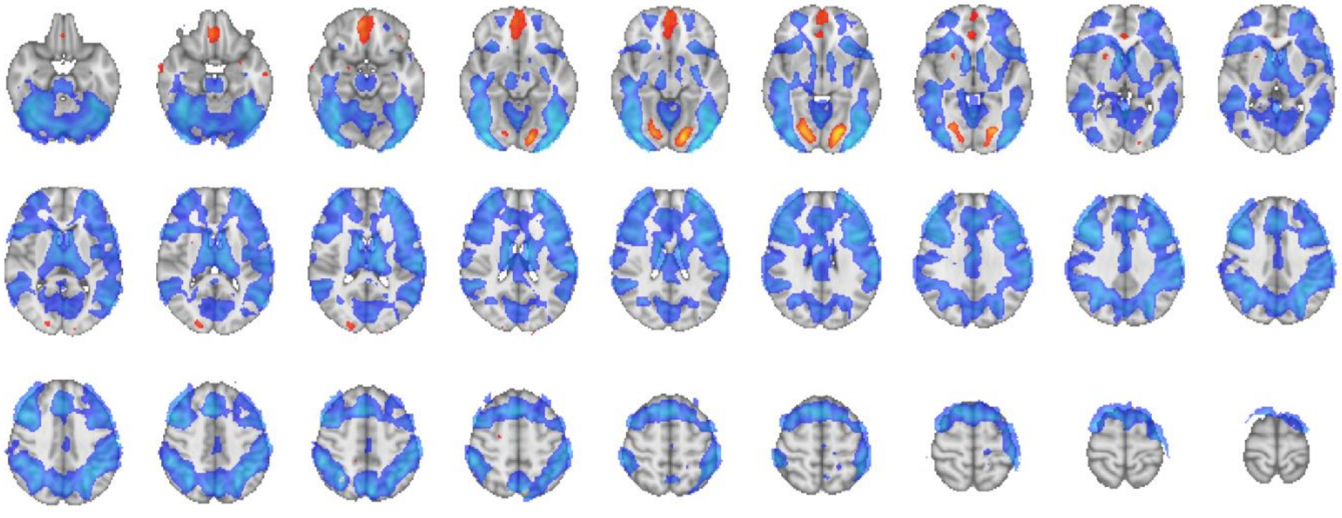

Feedback cue: Stand-alone fMRI version

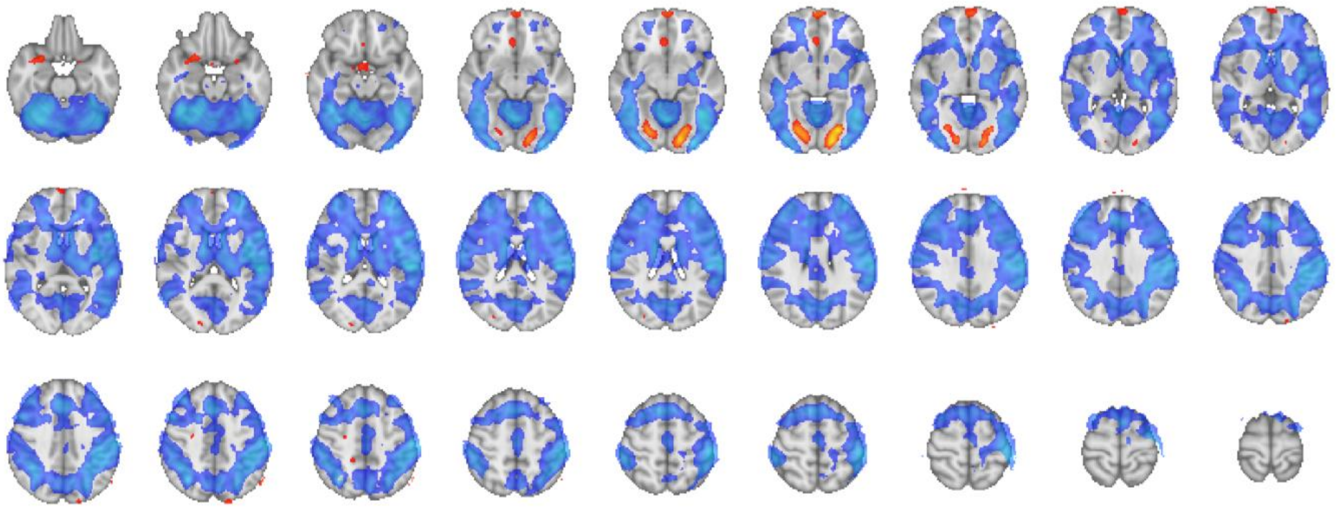

***Supplementary Figure 8.*** *Feedback Cue Z-statistics for the EEG-informed fMRI analysis and the stand-alone fMRI version. The stimulus onset regressor was a boxcar function with duration 0.1 and unmodulated amplitude. Red shows Z-statistics <= -2.57 and Blue, >= 2.57. No minimum cluster size was applied in this display.*

Explicit outcome value: EEG-informed fMRI Analysis

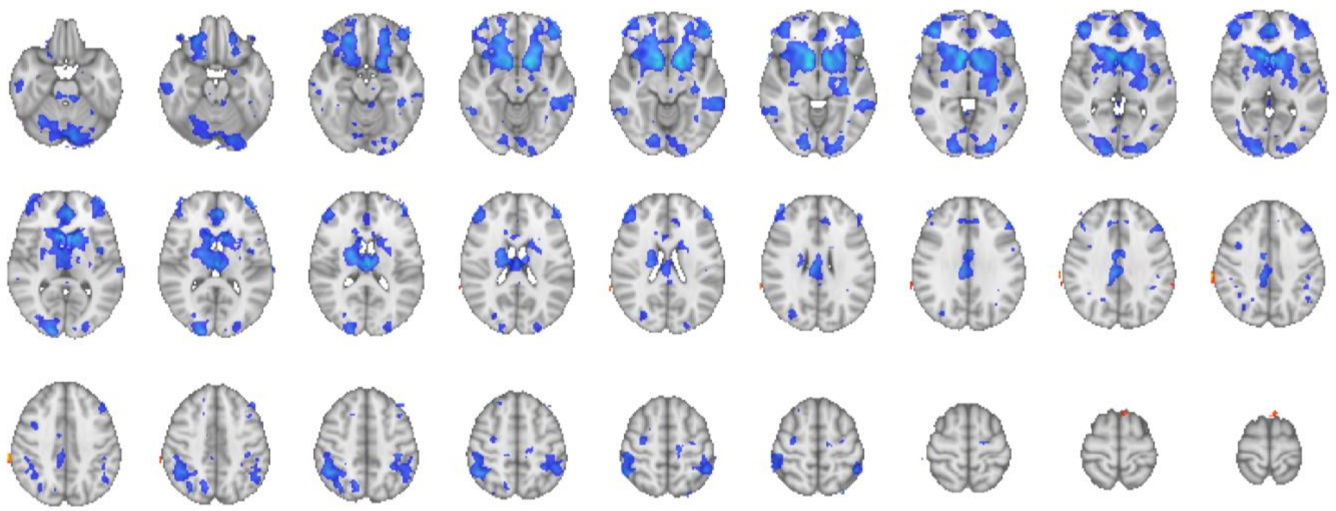

Explicit outcome value: Stand-alone fMRI version

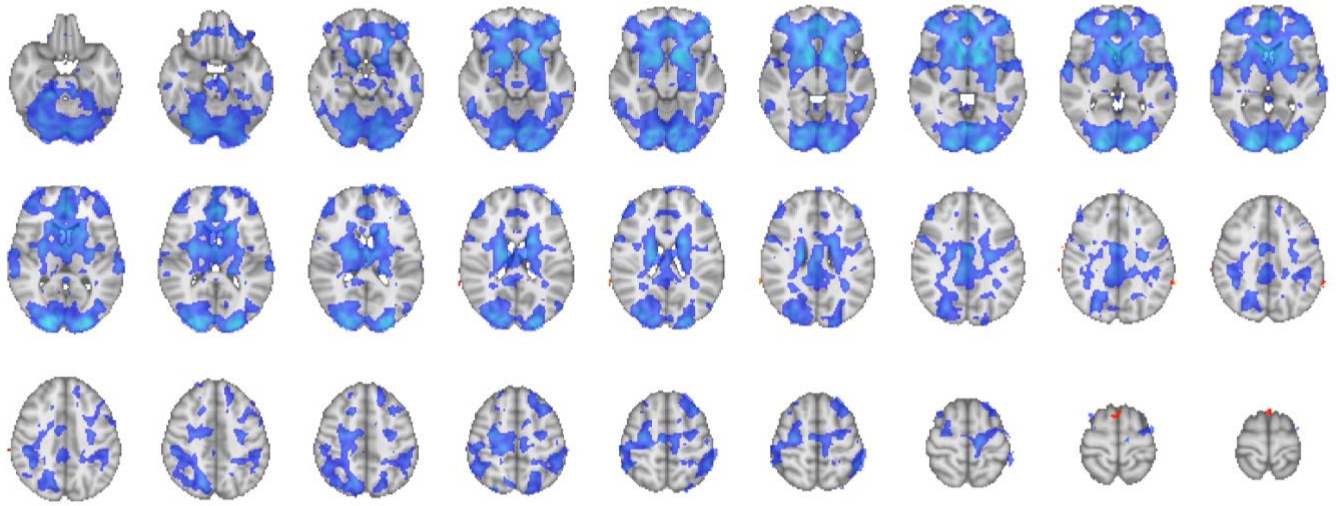

***Supplementary Figure 9.*** *Explicit outcome value Z-statistics for the EEG-informed fMRI analysis (EEG feedback-prediction following explicit feedback) and the stand-alone fMRI version (explicit feedback signed value). The regressor was a boxcar function with duration 0.1 and amplitude modulated by the EEG feedback-prediction following feedback (or the explicit feedback signed value for the stand-alone version). Red shows Z-statistics <= -2.57 and Blue, >= 2.57. No minimum cluster size was applied in this display.*

Implicit outcome value: EEG-informed fMRI Analysis

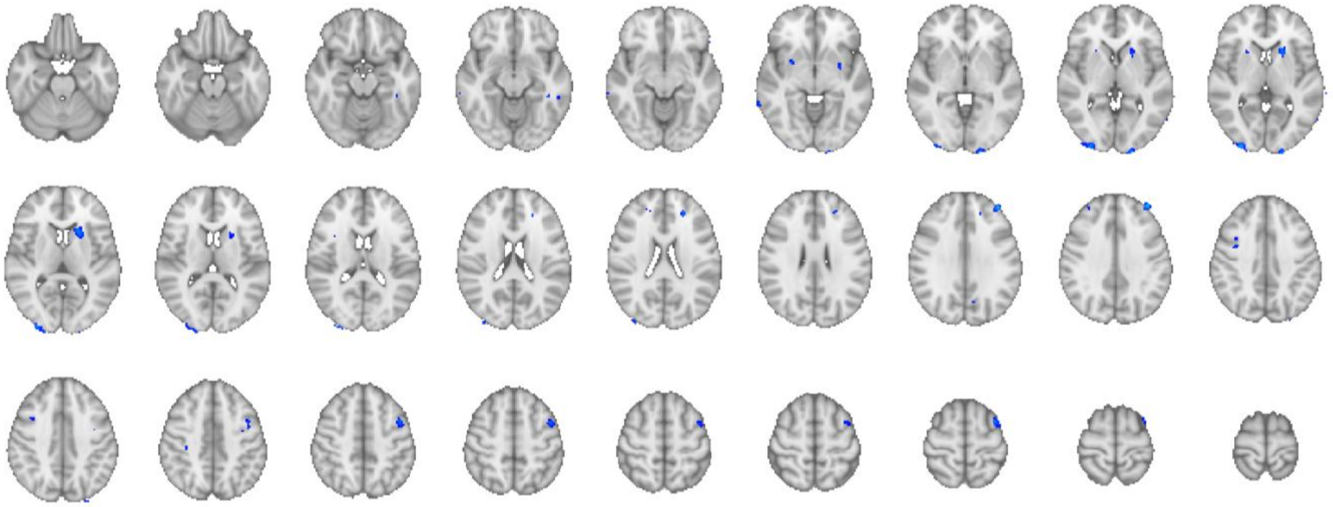

Implicit outcome value: Stand-alone fMRI version

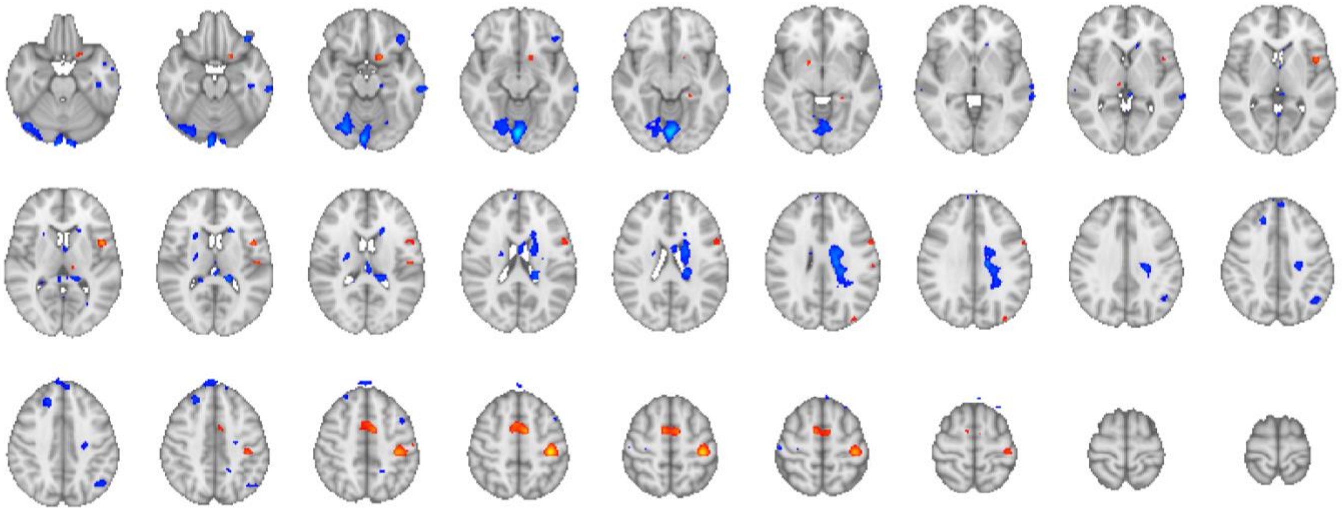

***Supplementary Figure 10.*** *Implicit outcome value Z-statistics for the EEG-informed fMRI analysis (EEG bet-prediction in the feedback-window, on no-feedback trials, top) and the stand-alone fMRI version (binary bet = 1, no-bet = -1 following no-feedback cue, bottom). The regressor was a boxcar function with duration 0.1 and amplitude modulated by the EEG bet-prediction following feedback (or the behavioural variable for the stand-alone version). Red shows Z-statistics <= -2.57 and Blue, >= 2.57. No minimum cluster size was applied in this display.*

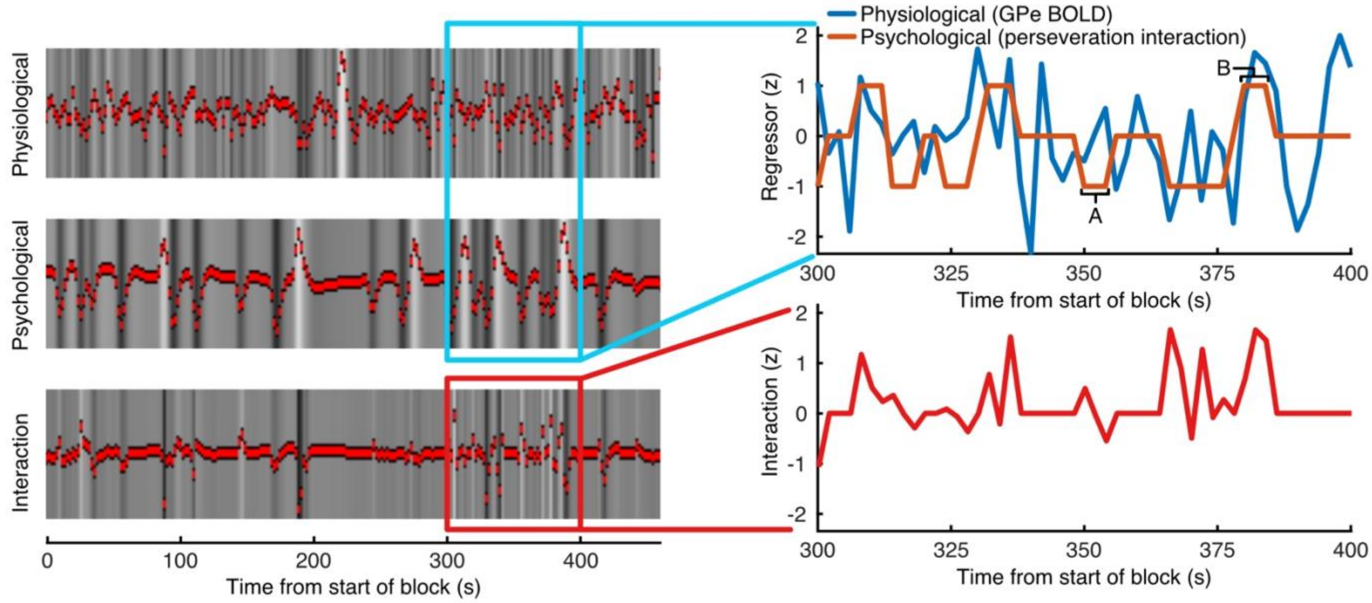


***Supplementary Figure 11. Psychophysiological interaction analysis design.*** *The images on the right show the resulting GLM regressors for one block from one subject. The physiological variable (top) is the GPe BOLD timeseries, taken as the average across voxels in the cluster. The psychological variable (middle) is the perseveration interaction, which is zero except in the time window from feedback over the intertrial interval, where it is set to 1 when the participant behaves according to learning from feedback (alternating responses to repeating stimuli following negative feedback (or no-bet responses on no-feedback trials), as well as trials with repeating responses to repeating stimuli following positive feedback (or bet responses on no-feedback trials), or -1 otherwise (repeating a response for a repeated stimulus following negative feedback (or no-bet trials), as well as alternating responses to repeating stimuli following positive feedback (or bet trials). The interaction is the variable of interest, where clusters of BOLD associated with this variable show increased connectivity with GPe when the participant's next response will be in line with learning from feedback (or decreased connectivity when the next response is not in line with learning from feedback). The panels to the right zoom in on the variables within the 300-400 s time-window. As a demonstration, the part of the psychological variable marked as A corresponds to the feedback window on trial 41 wherein the participant received negative feedback and on trial 42 the same stimulus direction was repeated and the participant repeated their response. The part of the psychological variable marked as B corresponds to the feedback window on trial 43 wherein the participant received negative feedback and on trial 44 the same stimulus direction was repeated but the participant changed their response.*

PPI: All trials

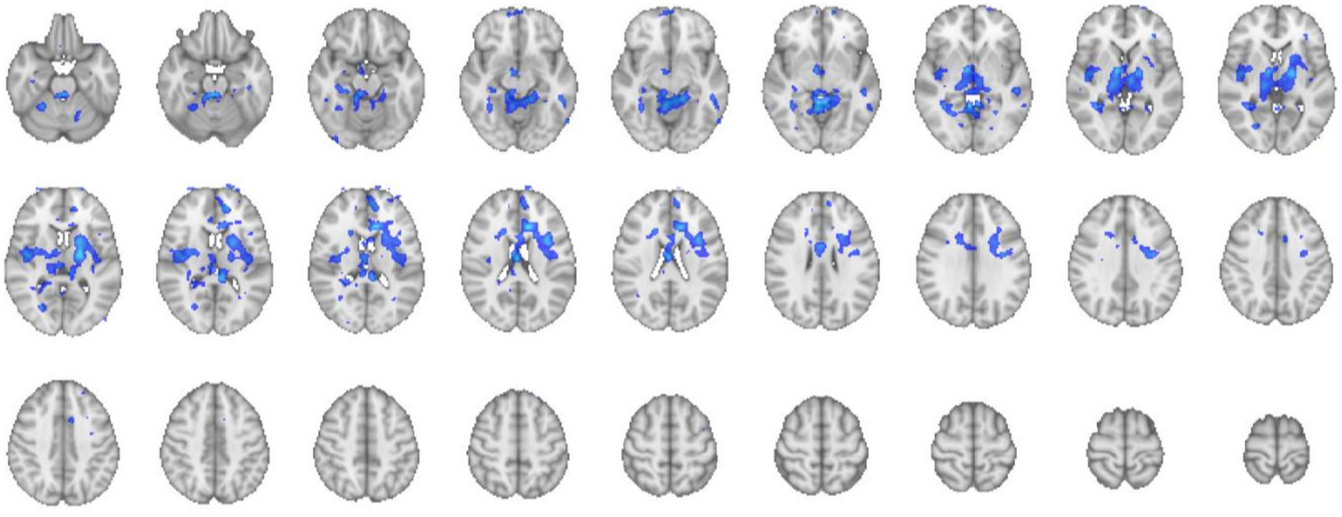

PPI: All trials vs. explicit-feedback trials

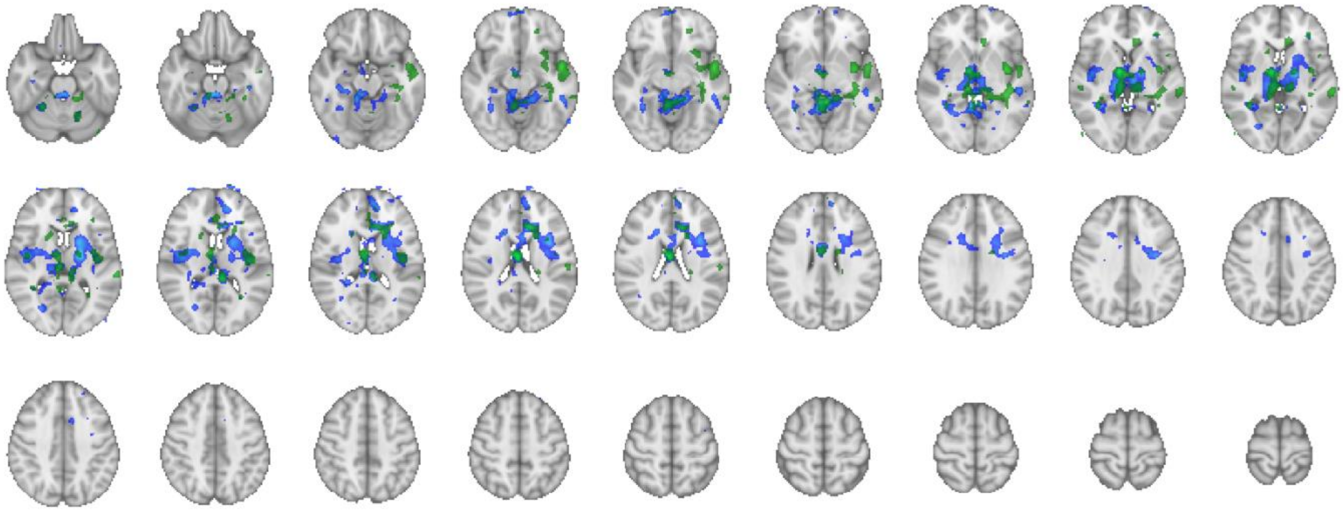

PPI: All trials vs. no-feedback trials

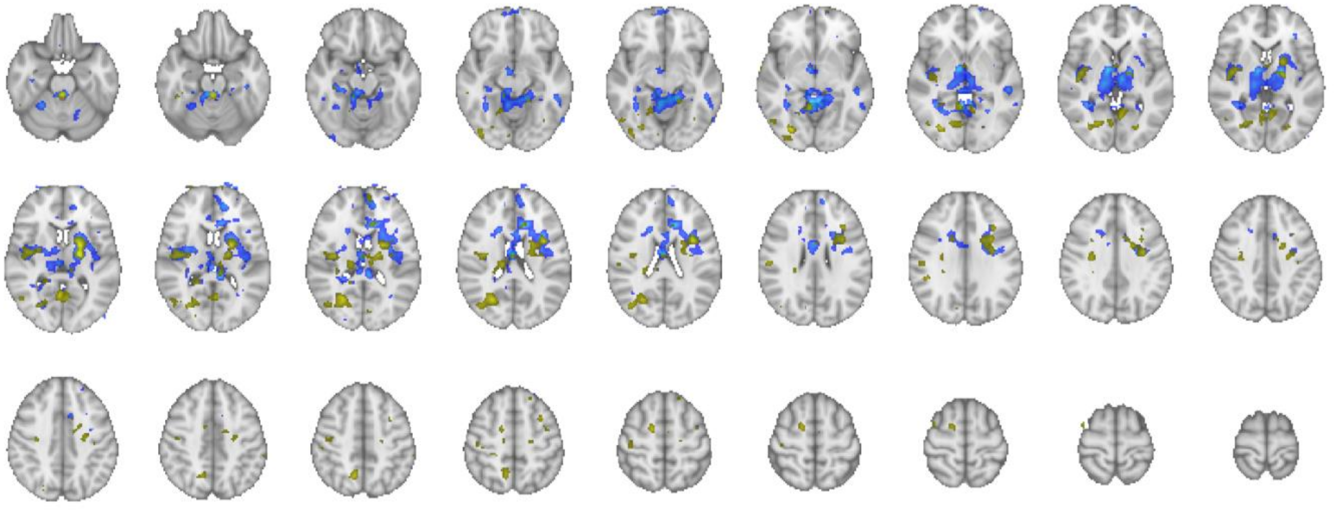

***Supplementary Figure 12.*** *Psychophysiological interaction Z-statistics for the analysis including all trials (top) compared to explicit-feedback trials (middle) and no-feedback trials (bottom). Coloured regions highlight Z >= 2.57 for the interaction between the BOLD time-course from the external globus pallidus region of interest and a variable coding the interaction between response perseveration and feedback (positive for repeating response following positive feedback and alternating response following negative feedback for repeating stimuli, the opposite combination for alternating stimuli, and otherwise negative).*

### Post-decision confidence

| Label | Voxels | MAX (Z-stat) | MAX X (MNI) | MAX Y (MNI) | MAX Z (MNI) |
|---|---|---|---|---|---|
| Left Motor/Somatosensory cortex including posterior middle frontal | 4407 | 5.72 | -34 | -24 | 60 |
| Left inferior frontal gyrus, GPe | 1558 | 4.13 | -60 | 8 | 22 |
| Right Parietal/somatosensory | 577 | 3.97 | 54 | -20 | 40 |
| Right inferior fronal gyrus / precentral gyrus | 510 | 4.23 | 54 | 4 | 36 |
| Left occipital | 388 | 4.41 | -40 | -66 | -8 |
| Cerebellum | 288 | 3.63 | 16 | -46 | -20 |
| Left rostrolateral PFC | 235 | 3.49 | -28 | 52 | 18 |
| Right middle frontal gyrus | 134 | 3.46 | 38 | 38 | 30 |

### Expected outcome value

| Label | Voxels | MAX (Z-stat) | MAX X (MNI) | MAX Y (MNI) | MAX Z (MNI) |
|---|---|---|---|---|---|
| Left occipital | 330 | 3.4 | -36 | -94 | -4 |
| Left parietal | 132 | 3.12 | -40 | -36 | 40 |
| Left insular | 124 | 3.79 | -50 | -20 | 20 |

### Explicit outcome value

| Label | Voxels | MAX (Z-stat) | MAX X (MNI) | MAX Y (MNI) | MAX Z (MNI) |
|---|---|---|---|---|---|
| Bilateral striatum, ventromedial and ventrolateral PFC, left Gpe | 14915 | 5.8 | -14 | -14 | -10 |
| Cerebellum | 6558 | 4.71 | -6 | -82 | -28 |
| Right Parietal | 1247 | 4.27 | 42 | -50 | 48 |
| Left superior parietal | 845 | 3.91 | -46 | -48 | 58 |
| Left middle temporal gyrus | 567 | 3.89 | -58 | -40 | -8 |
| Right middle temporal gyrus | 234 | 3.55 | 54 | -48 | -8 |
| Left Insular | 213 | 3.77 | -42 | 2 | 4 |
| Right temporal lobe | 188 | 3.49 | 56 | -24 | -20 |
| Bilateral paracingulate gyrus | 173 | 3.36 | 12 | 34 | 32 |
| Right Parietal | 156 | 3.73 | 14 | -66 | 46 |

### Implicit outcome value

| Label | Voxels | MAX (Z-stat) | MAX X (MNI) | MAX Y (MNI) | MAX Z (MNI) |
|---|---|---|---|---|---|
| Right occipital | 180 | 3.19 | 30 | -96 | 4 |
| Left middle frontal gyrus | 170 | 3.05 | -36 | 6 | 62 |
| Left dorsal striatum | 112 | 3.18 | -20 | 18 | 6 |

***Supplementary Table 1.*** *Cluster results for the fMRI GLM analysis (variables of interest only).*
